# Wnt/β-catenin inhibition disrupts drug-tolerance in isogenic carboplatin-resistant models of Triple-Negative Breast Cancer

**DOI:** 10.1101/2021.05.21.443409

**Authors:** Willy Antoni Abreu de Oliveira, Stijn Moens, Youssef El Laithy, Bernard K. van der Veer, Paraskevi Athanasouli, Emanuela Elsa Cortesi, Maria Francesca Baietti, Kian Peng Koh, Juan-Jose Ventura, Frédéric Amant, Daniela Annibali, Frederic Lluis

## Abstract

Triple-Negative Breast Cancer (TNBC) is the most aggressive breast cancer subtype, characterized by both limited treatment options and higher relapse rates than hormone-receptor-positive breast cancers. Chemotherapy remains the mainstay treatment for TNBC, and platinum salts have been explored as a therapeutic alternative in neo-adjuvant and metastatic settings. However, primary and acquired resistance to chemotherapy in general and platinum-based regimens specifically strongly hampers TNBC management. In this study, we used carboplatin-resistant in vivo patient-derived xenograft and isogenic TNBC cell-line models and detected enhanced Wnt/β-catenin activity correlating with an induced expression of stem cell markers in both resistant models. In accordance, the activation of canonical Wnt signaling in parental TNBC cell lines increases stem cell markers’ expression, formation of tumorspheres, and promotes carboplatin resistance. Finally, we prove that Wnt signaling inhibition resensitizes resistant models to carboplatin both in vitro and in vivo, suggesting the synergistic use of Wnt inhibitors and carboplatin as a therapeutic option in TNBC. Here we provide evidence for a prominent role of Wnt signaling in mediating resistance to carboplatin, and we establish that combinatorial targeting of Wnt signaling overcomes carboplatin resistance enhancing chemotherapeutic drug efficacy.

## 1 Introduction

Triple-negative breast cancer (TNBC) is a molecular subtype of breast cancer characterized by the lack of expression of estrogen-receptor, progesterone-receptor, and human epidermal growth factor receptor type 2 (1). TNBC accounts for 10-20% of all breast cancer cases, occurring with higher frequency in younger women, presenting with higher grade and mitotic counts than non-TNBCs, low differentiation, and frequent lymph node involvement, ultimately contributing to poor prognosis (1,2).

The lack of hormone- and growth factor-receptors render chemotherapy the primary systemic treatment for TNBC. Interestingly, TNBC patients have high response rates to neoadjuvant chemotherapy, achieving pathological complete response (pCR) more frequently than those bearing non-TNBCs (3). Nonetheless, TNBC patients experience lower progression-free- and overall survival rates and higher distant metastatic relapse frequency than non-TNBC patients, highlighting the critical need for alternative therapeutic approaches(3).

The Food and Drugs Administration of the United States of America first approved platinum salts, namely cisplatin, to treat metastatic testicular cancer, ovarian cancer, and bladder cancer between 1978 and 1979 (4). Since then, the use of platinum-based chemotherapy has grown and is now applied in many other cancer types. Pre-clinical studies have highlighted that TNBC is particularly sensitive to DNA damaging agents (5,6). For that reason, platinum salts – DNA-crosslinking agents – have gained traction as potential additions to the therapeutic toolbox for TNBC. Phase-II and Phase-III clinical trials have demonstrated the benefits of including carboplatin (CAR) in neoadjuvant regimens for TNBC (7–10). Importantly, pCR with neoadjuvant treatment is a robust predictor of survival in TNBC (3). However, chemotherapy-treated TNBC patients are likely to acquire resistance, and patients with residual disease (RD) have worsened prognosis and experience low survival rates, particularly within the first three years ensuing treatment (3).

During the last decades, our knowledge of platinum’s mechanism of action has increased significantly. Nonetheless, how cancer overcomes platinum-mediated cytotoxicity still holds unanswered questions. Several studies have shed light on how cancer cells adapt to platinum-based treatment by restoring DNA damage repair, increasing tolerance to DNA damage, decreasing its intracellular uptake and accumulation, and regulating apoptosis and autophagy (11). Chemotherapy resistance is also known to be induced and maintained by adaptations in pro-survival and anti-apoptotic signaling pathways. Like other chemotherapeutic agents, alterations of such cellular dynamics also affect platinum-based treatments. Several studies have demonstrated the involvement of NOTCH (12), MEK (13), Hedgehog (14), EGFR (15), among others, in mediating resistance to platinum in different cancer types. Also, cancer cells with stem cell-like properties have been described to significantly influence the response to different chemotherapeutic agents, including platinum compounds (16–19). Cancer stem cells constitute a subpopulation of cancer cells with tumorigenic and self-renewal capacities and are considered desirable therapeutic targets since their intrinsic cellular properties contribute extensively to treatment failure (20–22). Breast cancer stem cells were first isolated in 2003 based on cell surface markers CD44 and CD24 (23). Since then, many studies have demonstrated their tumorigenic and drug-resistance capacities, highlighting the need to develop therapeutic approaches that deplete this population (6,24–26).

The Wnt/β-catenin signaling pathway is a developmental signaling cascade with a prominent role in cancer (27). It is initiated when Wnt ligands (secreted lipid-modified signaling molecules) bind the receptor complex at the cell membrane. A series of events ensues, culminating in the inhibition of glycogen synthase kinase 3 beta (GSK3β), and the subsequent cytoplasmic accumulation of β-catenin, the key mediator protein of Wnt signaling. This accumulation leads to the nuclear translocation of β-catenin eliciting Wnt target genes’ expression by interacting with different transcription factors. In the absence of Wnt ligands, β-catenin is constitutively phosphorylated by GSK3β and targeted for proteasomal degradation (28). Notably, Wnt is known to govern several cellular functions with the potential to contribute to chemotherapy resistance. Such functions include the control and regulation of proliferation (29), DNA damage repair (30), inhibition of apoptosis (31), and maintenance and regulation of embryonic, somatic, and cancer stem cell properties (32). Different studies have demonstrated Wnt pathway involvement in the mediation of platinum resistance in various cancer types, including squamous cell carcinoma (33,34) and ovarian cancer (35). However, its involvement in platinum resistance in TNBC is not known.

To study how TNBCs acquire resistance to carboplatin treatment, we used a carboplatin-tolerant isogenic TNBC cell line. Transcriptomic analysis was performed to gain insight into biological signaling pathways underlying acquired carboplatin resistance *in vitro*, leading to the identification of Wnt signaling as a candidate resistant-mediating pathway. Additionally, the resistant cell line displayed enhanced expression of pluripotency markers and stem cell features compared to the parental, carboplatin-sensitive cells.

*In vitro* pharmacological and genetic manipulation of Wnt signaling was employed to assess drug response alterations and stem cell potential functionally. Inducing Wnt signaling in parental non-resistant TNBC cell lines elicited the expression of pluripotency markers observed in isogenic resistant cells and enhanced stem cell features *in vitro*. Moreover, pharmacological and genetic inhibition of Wnt activity in resistant cells disrupted carboplatin tolerance and hindered tumorsphere formation. Finally, carboplatin-tolerant isogenic patient-derived xenograft (PDX) models were used to test the effect of *in vivo* Wnt inhibition on platinum-response. Similar to what we observed *in vitro*, inhibition of Wnt reduced expression of cancer stem cell markers and drastically reduced tolerance to carboplatin treatment *in vivo*.

Altogether, our results suggest the potential for Wnt signaling inhibition in combination with carboplatin as a strategy to prevent or overcome platinum resistance in TNBC patients.

## 2 Materials and methods

### Cell lines, cell culture, and treatments

MDA-MB-468 (ATCC-HTB-132) and MDA-MB-231 (ATCC HTB-26) were maintained in DMEM high glucose (Gibco 41965039) supplemented with 10% fetal bovine serum, 1mM sodium pyruvate (Gibco, 11360070), 1X non-essential amino acids (Gibco, 11140035), 100μg/mL penicillin-streptomycin (Gibco, 15140163) and 0.01mM 2-mercaptoethanol (Gibco, 31350010).

Unless otherwise specified in the text, all carboplatin treatments (CAR, Hospira UK, Ltd) were done at 2μM for MDA-MB-468 cells and 35μM for MDA-MB-231 cells. Small molecule Wnt activator CHIR99021 (CHIR, Sigma, SML1046) was administered to cells in DMSO (Sigma, D2650) at 4μM. Small molecule Wnt inhibitor LGK-974 (LGK, Selleckchem, S7143) was, unless otherwise specified, administered at 200 nM in DMSO.

Carboplatin resistance was induced in MDA-MB-468 cells by continued maintenance in carboplatin containing medium, starting at a concentration of 0.4 μM. The concentration of carboplatin was increased in 9 increments until reaching 2 μM, once unhindered cell growth was obtained at each concentration level, allowing a 48h carboplatin-free recovery period with each splitting (35).

### Lentiviral particle production and transduction

Lentiviruses were produced according to the RNAi Consortium (TRC) protocol available from the Broad Institute (https://portals.broadinstitute.org/gpp/public/resources/protocols). In brief, 5×10^5^ HEK293T cells seeded per well in 6-well plates and transfected the following day with 750 μg pCMV-dR8.91, 250 μg pCMV-VSV-G, and 1 μg of the specific lentiviral expression or silencing constructs using FugeneHD (Promega, E2311) in Opti-MEM (Gibco, 31985070). One day after, the culture medium was replaced. The same day, lentivirus-recipient cells were plated in 6-well plates at a density of 5×10^4^ cells per well. Lentivirus-containing medium was collected from HEK293T cells 48h and 72h after transfection and added to recipient cancer cells after being filtered. Two days after infection, cells were washed thoroughly with PBS, medium refreshed, and appropriate selection antibiotics applied.

For overexpression of ΔN90-β-Catenin, we used pLV-beta-catenin ΔN90 (Addgene, #36985) and pPRIME-CMV-NEO-recipient (CTRL, Addgene, #11659). For β-Catenin shRNA mediated silencing, we used pXL002-ishRNA-beta-catenin-1 (Addgene, #36297) and pXL004-ishRNA-scramble (Addgene, #36311). For Wnt fluorescent reporter assay, we used 7TGP (Addgene, #24305).

### In vitro carboplatin response (IC50)

For IC50 experiments, we seeded 2.5×10^4^ cells per well in 12-well plates. Cells were treated with increasing concentrations of CAR (0.02 to 200 μM for MDA-MB-468 and 0.35 to 3500 μM for MDA-MB-231) for 72 hours. Viability was assessed by manual cell counting using a Neubauer hemocytometer using trypan blue for dead cell exclusion. Cell viability was determined as a percentage of untreated cells, and non-linear regressions of [CAR] vs. normalized-response were fitted using GraphPad Prism v.8.0.1. to mathematically determine the IC50.

### Flow Cytometry

For annexin V apoptosis analysis, cells were detached and resuspended in annexin V binding buffer (BD Pharmigen, 51-66121E) and incubated for 15 minutes at room temperature with APC-conjugated AnnexinV (Thermo-eBioscience, BMS306APC-100). After incubation, cells were diluted in binding buffer containing 100 nM of 4’,6-diamidino-2-phenylindole (DAPI). Unstained and single-stained (annexin V and DAPI) were used as gating controls.

For ALDH activity assays, cells were detached, washed in PBS, and stained using the AldeRed ALDH detection assay kit (Merck SRC150) according to manufacturer specifications.

For immunolabeling of CD44 and CD24, cells were detached, washed twice in PBS with 4% FBS, and incubated with CD44-PE (BD Pharmigen, 555479) and CD24-APC (Invitrogen, 17-4714-81) antibodies according to manufacturer specifications at room temperature. After incubations, cells were washed twice in PBS with FBS and resuspended in PBS containing 4% FBS and 100 nM of DAPI. Cells incubated with PE- and APC-conjugated isotype-antibodies and single-stained cells were used as gating controls.

All data were collected on a BD FACS Canto II at the KU Leuven Flow Cytometry Core and analyzed using FlowJo v.10.6.2.

### SDS-PAGE and Western Blot

For western blot, cells were collected and washed in PBS before being pelleted. Then, cells were lysed on ice with RIPA buffer (150 mM NaCl, 1% Nonidet P40, 0.5% sodium deoxycholate, 0,1% dodecyl sulfate, 50 mM Tris-HCL, pH 8.0) containing a cocktail of protease and phosphatase inhibitors (Sigma, #P5726, #P0044, #P8340). Lysates were centrifuged at 16.000x g for 10 minutes at 4°C to discard insoluble material, and protein concentration was determined using the Bradford method. For SDS-PAGE, 30 μg of protein were mixed with 4x Laemmli buffer (240 mM Tris/HCL pH 6.8, 8% SDS, 0.04% bromophenol blue, 5% 2-mercaptoethanol, 40% glycerol) and denatured for 5 minutes at 96°C prior to electrophoretic protein separation. Resolved protein extracts were transferred to PVDF membranes (BIORAD, 162-0177). Transfer success was assessed with Ponceau S solution, and membranes were blocked with 5% non-fat milk or 5% BSA in TBS-T (0,1% Tween-20^®^) for 60 minutes. After blocking, membranes were incubated with primary antibodies at 4°C overnight. The day after, membranes were washed 3 times with PBS-T for 10 minutes and incubated with secondary HRP-conjugated antibodies. Immunolabeled proteins were detected with Supersignal West Pico chemiluminescent kit (Fisher Scientific, 34077) on autoradiography film (Santa Cruz, SC-201697). The primary antibodies used were active rabbit anti-non-phosphorylated β-Catenin (CellSignaling Technologies, #19807S), mouse anti-total β-Catenin (BD, #610154), mouse anti-β-actin (Santa Cruz, #47778).

### Next-generation mRNA Sequencing

For mRNA sequencing, total RNA was obtained from cells using the GenElute mammalian total RNA miniprep kit (Sigma, RTN350-1KT). Libraries were prepared from 250 ng of total RNA using the KAPA stranded mRNA-seq kit (Roche, KK8421) according to the manufacturer’s specifications. KAPA-single index adapters (Roche, KK8700) were added to A-tailed cDNA, and libraries were amplified by 12 cycles of PCR. Finally, libraries were purified on Agencourt AMPure XP beads (Beckman Coulter, A63881). Libraries were controlled for fragment size using the High Sensitivity DNA analysis kit (Agilent, 5067-4626) on an Agilent Bioanalyzer 2100. Each library was diluted to 4 nM and pooled for single-end 50-bp sequencing on an Illumina Hiseq4000 20 – 27 million reads per sample (22 million reads on average).

Adapters, polyA tails, and bad quality reads (Phred score > 20) were trimmed using Trim Galore! (v0.6.4_dev) with default parameters. Reads were aligned to the transcriptome and quantified using Salmon (v0.14.1) (36) with default parameters using GENCODE release 36 of the human reference transcriptome sequences and the comprehensive gene annotation. Subsequently, the counts were imported into R (v4.0.2) using tximport (v1.18.0) and differentially expressed genes were defined using DEseq2 (v1.30.0) (37)and log fold changes corrected using “ashr” method (38) (FDR adjusted p.val < 0.05 & |log2(fold change)| > 1.5). TPM values were also calculated using tximport.

### Functional enrichment analysis and enrichment maps

Datasets GSE103668 and E-MTAB-7083 were downloaded from the GeneExpression Omnibus and ArrayExpress public repositories, respectively. Differentially expressed genes with |log2(fold-change)| > 1 and p-value < 0.05 were obtained using limma (v3.26.8) R package in R (v4.02) and by using the limma method on NetworkAnalyst (39). Differentially expressed genes were ranked by fold-change for Gene Set Enrichment Analysis (GSEA v4.1.0) using weighted enrichment statistic and KEGG, Hallmarks, and Wikipathways gene sets. Additionally, we used custom gene sets comprised of human embryonic stem cell-related genes (M1871: BENPORATH_ES1 and M4241: BENPORATH_ES2), pluripotency transcription factor target genes (M14573: BENPORATH_NOS_TARGETS), and cancer progenitor genes (ENGELMANN_CANCER_PROGENITORS_UP) obtained from www.gsea-msigdb.org. The statistical signifcance threshold was set at FDR<0,1 or (*p*<0,5 **∧** FDR<0,25). Additionally, gProfiler (https://biit.cs.ut.ee/gprofiler/gost) was used to assess the function of ranked DEGs using the ranked query mode and Benjamini-Hochberg FDR thresholding. The outputs of GSEA and gProfiler analysis were fed to the EnrichmentMap app on Cytoscape (v3.8.1) to generate visualizations of enriched biological features and pathways following published protocols (40). Differentially expressed genes from RNA-sequencing were processed for functional analysis and visualization in the same way, except for GSEA, differentially expressed genes were ranked by the absolute value of fold change (41).

### Real-Time Quantitative Polymerase Chain Reaction and gene expression analysis

For RT-qPCR, total RNA was extracted using the GenElute mammalian total RNA miniprep kit from Sigma (Sigma, RTN350-1KT) according to the manufacturer’s instructions, and DNA was digested during RNA extraction using on-column DNAse (Sigma, On-Column DNAse I digestion set, DNASE70). cDNA was synthesized from 500 ng of total RNA using the BIORAD iScript cDNA synthesis kit (BIORAD, CAT#1708891). Quantitative real-time PCR reactions were set up in technical triplicates with Platinum SYBR Green qPCR SuperMix-UDG (Invitrogen, 11733-046) on a ViiA7 Real-Time PCR System (Thermo Scientific). Expression levels were normalized to housekeeping genes (HKG) GAPDH and RPL19. Statistical testing of differences in expression between samples was carried out on relative-expression values (2^−ΔCT^). In some figures, mRNA expression values are represented as fold-change for convenience of interpretation, although statistical testing was performed on relative expression values (2^−ΔCT^).

### Tummorsphere formation assays

For tumorsphere formation assays, cells were collected as described above, washed, counted, and resuspended in serum-free tumorsphere assay medium containing DMEM/F12, 1x B27 (Thermo, 12587010), 10ng/mL bFGF, (Peprotech, 100-18b) 20 ng/mL EGF (Peprotech, AF-100-15), and 2% growth-factor reduced matrigel (Corning, 734-0268). Cells were seeded at a density of 1000 cells/mL in ultra-low attachment 6-well plates and allowed seven days to grow. On the 7^th^ day, spheres were collected and centrifuged at 50g for 10 minutes, resuspended, and transferred to 96-well plates. Plates were briefly centrifuged at 50g for 1 minute to pull down larger spheroids (>60 μm) which were counted using a tally counter.

### Immunohistochemistry

Tumor samples were dissected, washed in saline, and either snap-frozen in OCT compound (VWR 361603E) or fixed in 4% formalin for 24 hours. Frozen tissue was cut at 10 μm thickness using a cryostat and mounted on superfrost microscope slides (Thermo Scientific, J1800AMNZ). Formalin-fixed tissue was embedded in paraffin and sectioned at 4 μm thickness using a microtome. Frozen and FFPE sections were stained with hematoxylin and eosin (HE), using a Leica Autostainer XL (Leica Microsystems), or stained in immunohistochemistry (IHC). In brief, frozen sections were fixed in acetone and preserved at −80°C until use. Slides were thawed at room temperature for 15 minutes and rehydrated in PBS. FFPE sections were deparaffinized in Leica Autostainer XL (Leica Microsystems) and pre-treated in citrate buffer (EnVision FLEX Target Retrieval Solution Low pH, Agilent-Dako, K8005) using a PT Link module (Dako), according to manufacturer’s instructions. For IHC, tumor sections were incubated with Envision Flex Peroxidase-Blocking Reagent (Dako, S202386-2) for at least 5 minutes, rinsed three times in wash buffer (Dako, K800721-2), and blocked with 5% bovine serum albumin (BSA) for 45 min at room temperature. After overnight incubation with anti-human Ki67 antibody (Abcam, EPR3610 clone, 1:1500) at 4°C, slides were incubated with HPR-conjugated secondary antibodies (Agilent, K400311-2) for 30 min. Tissue sections were stained with 3,3’-diaminobenzidine solution (DAB; Liquid DAB+ Substrate Chromogen System, K346889-2, Dako), counterstained with hematoxylin, and mounted using an automated coverslipper machine (Leica CV5030, Leica Biosystems). Pictures were acquired using a Zeiss Axiovision microscope. Quantification of Ki67+ cells was performed in at least 5 random 20x fields per sample using QuPath 0.2.3. (42).

### Tunel Staining and pan-cytokeratin immunofluorescent staining

For TUNEL staining, cryopreserved tumor samples were cryosectioned at 10 μM thick and mounted on superfrost microscope slides. Slides were stored at −80°C and stained using the Click-iT Plus TUNEL (ThermoFisher C10617) according to the manufacturer’s instructions. After TUNEL staining, slides were blocked with 5% normal donkey serum in PBS (Gibco, 10010-050) for one hour and incubated overnight at 4°C with rabbit anti-pan-cytokeratin polyclonal antibody (Abcam ab217916 1:400). The following day, slides were washed three times in PBS containing 0.01% Triton X-100 and incubated for 2 hours with AlexaFluor conjugated donkey anti-rabbit secondary antibody (Abcam ab150073 1:1000). After secondary antibody incubation, slides were washed three times with PBS containing 0.01% Triton X-100 and mounted with ProLong™ Gold Antifade Mountant with DAPI (P36931, Thermo Fisher Scientific). Images were acquired using a Leica Sp8x confocal microscope. Quantification of TUNEL positive/ pan-cytokeratin positive cells was done in at least eight randomly sampled 10x fields per sample using QuPath 0.2.3.

### PDX models

BRC016 (primary, grade III, TNBC) was established at the University Hospital UZ Leuven and is available from the Trace Leuven Cancer Institute (https://www.uzleuven-kuleuven.be/lki/trace/trace-leuven-pdx-platform). C4O was previously obtained from a carboplatin treatment-refractory BRC016 tumor. The regrown tumor was harvested and implanted on NMRI-Fox1nu nude mice (Taconic) for propagation, re-testing, and confirmation of carboplatin tolerance in a previously published study (35). Treatment experiments included 24 NMRI-Fox1nu nude mice implanted with C4O tumor fragments. When tumors reached a volume of approximately 300 mm^3,^ mice were randomly assigned to placebo (vehicle), CAR (50 mg/kg), LGK (2 mg/kg), or CAR+LGK (50 mg/kg + 2 mg/kg) treatment groups. Carboplatin was administered once a week intraperitoneally, and LGK974 was administered daily by oral gavage. Treatments were carried out for three weeks. Tumors were measured every 48 hours with digital calipers, and volume was estimated as V = L × W^2^ x π/6 (L: length, W: width).

### Statistical analysis

All data were analyzed using GraphPad Prism 8, except for transcriptomic datasets. Unless otherwise specified, comparisons between two groups were tested for statistical significance using unpaired t-tests with Welch correction. Comparisons between multiple groups were performed using a one-way analysis of variance (ANOVA). Comparisons between multiple groups across multiple time points were performed using two-way ANOVA. All statistical testing was corrected for multiple comparisons, using the Holm-Sidak method when comparing samples based on experimental design or the Tukey method when testing the comparison between all means in a dataset. For the reader’s convenience, all statistical tests and sample sizes are indicated in the figure legends.

## 3 Results

### Carboplatin-tolerant TNBC cells are characterized by enhanced WNT/β-catenin pathway activity, stem cell marker expression, and tumorsphere formation capacity

To explore the mechanisms of *in vitro* carboplatin resistance in TNBC, we generated an isogenic carboplatin-tolerant cell line (468’CT) (Fig. 1A) by exposing MDA-MB-468 cells to carboplatin treatment in incremental cycles.

**Figure 1.**
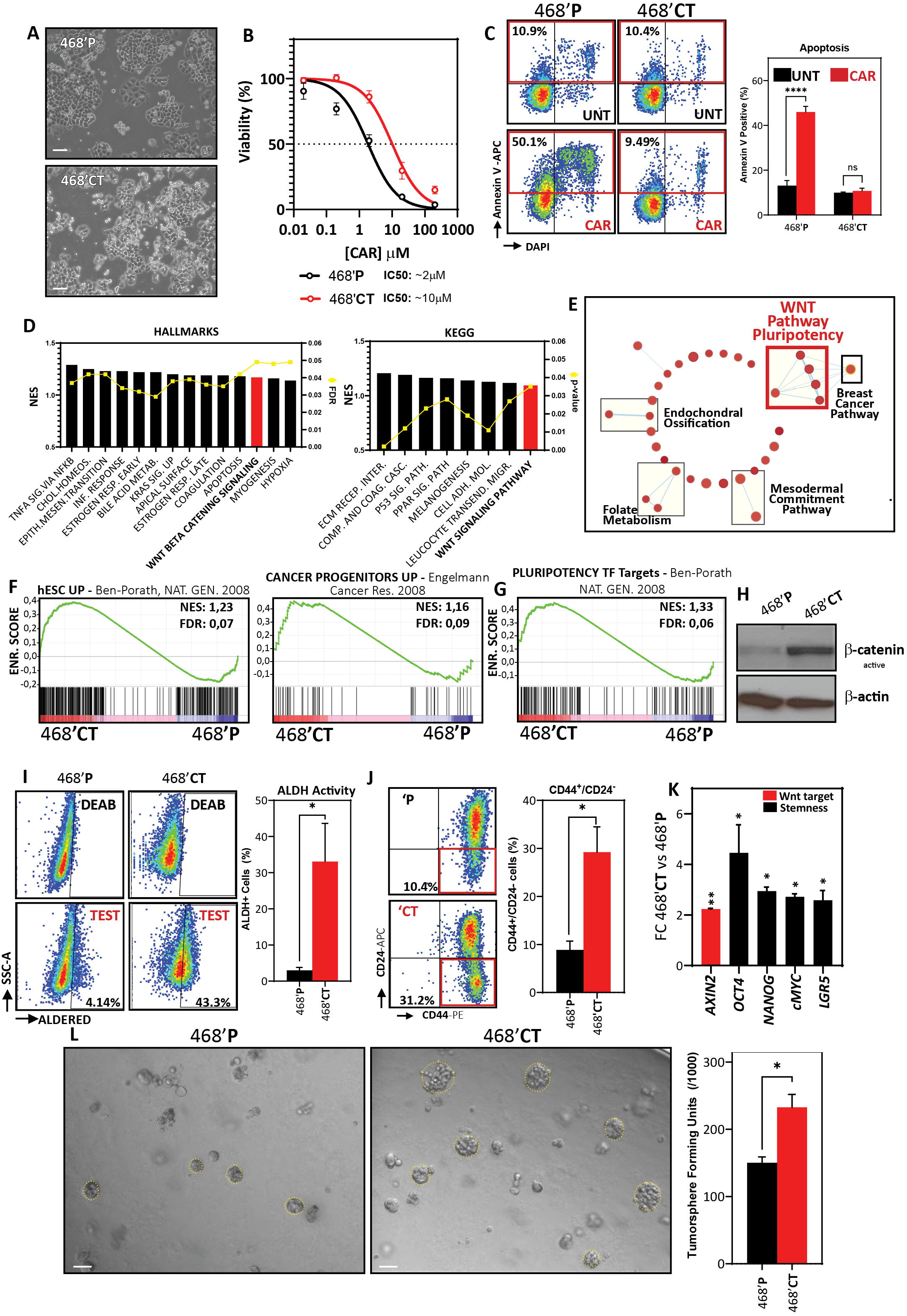
Carboplatin-tolerant TNBC cells are characterized by enhanced WNT/β-catenin pathway activity, stem cell marker expression, and tumorsphere formation capacity. A) Phase contrast microscope images of 468’P and 468’CT cells. Scale bar: 100 μm. B) Non-linear fit model of [CAR] μM vs normalized response for IC50 determination. (468’P: n=6, R^2^=0.92) (468’CT: n=4, R^2^=0.95). C) Representative flow cytometry scatterplots of annexin V staining of cells treated with 2 μM CAR for 72 hours (left) and respective statistical analysis (right) using multiple t-tests corrected for multiple comparisons with the Holms-Sidak method (n=3). D) Enriched gene sets from Hallmarks and KEGG databases by one-tailed GSEA ranked by Normalized enrichment score (NES), illustrating pathways most significantly deregulated between 468’P and 468’CT. E) Enrichment map of one-tailed GSEA hits from Wikipathways database. Rectangles highlight clusters of gene sets with significant overlap and are labeled using AutoAnnotate on Cytoscape. F) GSEA of hESCs (26)(left) and Cancer Progenitor(44) (right) gene sets in 468’CT vs 468’P cells. G) GSEA of NANOG, OCT4 and SOX2 target genes determined by ChIP-SEQ in hESC’s (26) in 468’CT vs 468’P cells. H) Western blot of active non-phosphorylated β-catenin in 468’P and 468’CT. β-actin was used as the loading control. I) Representative scatterplots of flow cytometric analysis of aldehyde dehydrogenase activity (left). DEAB panels refer to internal controls in which ALDH activity is blocked with diethylaminobenzaldehyde to determine background signal generated by unconverted ALDH substrate. TEST panels refer to the experimental samples where substrate for fluorimetric determination of ALDH activity is supplied. TEST samples are normalized to background fluorescence measured in DEAB internal controls and presented as the mean + standard error of the mean percentage of ALDH+ cells in 468’CT (n=5) and 468’P (n=7) (right). Welch’s t-test. J) Representative scatterplots of flow cytometric analysis of CD44-PE and CD24-APC immunolabeling (left) and corresponding statistical analysis of the mean percentage of CD44^+/^CD24^−^ cells (right; n=3). Welch’s t-test. K) qRT-PCR of Wnt target AXIN2 and stem cell markers in 468’CT cells vs. 468’P (n=4). Multiple t-tests. L) Representative brightfield images of tumorspheres generated from 468’P and 468’CT cells (left, scale bar: 50 μm) and statistical analysis of mean tumorsphere forming units (number of spheres/number of seeded single cells) (right; n=3). Welch’s t-test. (Barplots represent mean + SEM. * *p <0.05, ** p<0.01, *** p<0.001, **** p<0,0001*, ns= non significant).

Half-maximal inhibitory concentration (IC50) profiles for carboplatin were determined for both parental (468’P) and 468’CT cells (Fig.1B). The carboplatin-tolerant cell line displayed a 5x increase in IC50, thereby functionally confirming a significant increase in carboplatin tolerance. Flow-cytometric analysis of apoptosis further corroborated the establishment of the carboplatin-tolerant phenotype. When exposed to 2μM for 72 hours, 468’P stained positively and significantly for the apoptosis marker annexin V (43), whereas no significant increase in apoptotic cells was observed in 468’CT (Fig. 1C).

To study the underlying mechanisms of carboplatin tolerance, we performed transcriptome analysis by next-generation mRNA sequencing in 468’P and 468’CT cells. We used the ranked differentially expressed genes (Supplementary Table 1) followed by one-tailed Gene Set Enrichment Analysis (GSEA) (41) to identify changes in signaling pathways (Hallmarks, KEGG and Wikipathways). GSEA analysis identified alterations in several key cancer-related processes such as epithelial-mesenchymal-transition (Hallmarks) and PPAR and P53 signaling (KEGG) (Supplementary Table 1). However, Wnt signaling was consistently enriched across the two databases (Fig. 1D) with a clear differential expression pattern across the two cell lines (Supplementary Fig. 1A). Moreover, enrichment maps of Wikipathway database terms highlighted a cluster of gene sets comprising Wnt signaling and pluripotency regulation (Fig. 1E), suggesting a potential acquisition or enrichment of stem cell features in carboplatin tolerant cells.

To further understand the differences in stem cell transcriptional features between 468’CT and 468’Pcells, we compared our transcriptomic data with a curated gene sets comprised of genes found overexpressed in human embryonic stem cells (hESC) and cancer stem cells (26,44). Interestingly, 468’CT seem to be transcriptionally closer to both embryonic and cancer stem cells than 468’P as determined by GSEA (Fig. 1F). In addition, we compared our transcriptomic data with a gene set comprised of targets of pluripotency transcription factors NANOG, OCT4, and SOX2 in hESCs determined by chromatin immunoprecipitation followed by DNA sequencing (26). GSEA revealed enrichment of pluripotency transcription factor target genes in 468’CT cells, lending further support to the putative acquisition of stem cell features in this cell line (Fig. 1G).

To functionally validate the observed differences in Wnt/β-catenin signaling between 468’CT and 468’P cells, we analyzed protein levels of non-phosphorylated (active) β-catenin. Importantly, western-blot analysis of total protein extracts in baseline untreated conditions revealed strong enrichment of active β-catenin in 468’CT cells compared to the parental counterpart, thereby confirming the functional activation of Wnt activity in drug-tolerant cells (Fig. 1H). Moreover, the accumulation of active β-catenin was accompanied by transcriptional activation of Wnt-reporter activity (Supplementary Fig. 1B).

To quantify differences in frequency of putative cancer stem cell populations in both cell lines, we used flow cytometry to assess the enzymatic activity of aldehyde dehydrogenases (ALDH) and the expression level of the cell surface markers CD44 and CD24. Both methods have been used to identify, quantify, and isolate putative cancer stem cells from different cancer types. High ALDH activity and CD44/CD24 expression ratio in TNBC have been shown to correlate with enhanced tumorigenesis and metastatic potential as wells as radio- and chemotherapy resistance (23,25,45). Flow cytometric analysis showed significant differences in ALDH positive (Fig. 1I) and CD44^+^/CD24^−^ cells (Fig. 1J), with 468’CT cells expressing higher levels of both markers. Gene expression analysis also revealed an enrichment of the core pluripotency regulators *OCT4, NANOG*, and *cMYC*, as well as cancer stem cell marker *LGR5* (46) (Fig. 1K).

To functionally evaluate differences in cancer stem cell properties and *in vitro* tumor-initiating capacity, we performed a tumorsphere formation assay. *In vitro* growth in non-adherent conditions has been described as an exclusive capability of cancer stem cells, thereby functioning as a surrogate measure of *in vitro* tumor-initiating capacity and as a method to enrich cancer stem cells (47). Importantly, in line with our stemness-related gene expression data and flow cytometry analysis of ALDH activity and CD44/CD24 expression, we observed a significantly higher tumorsphere formation frequency in 468’CT cells compared to 468’P (Fig. 1L).

Altogether, the transcriptomic evidence for alterations in Wnt signaling, and presumably stem cell features, between 468’CT and 468’P cells suggests the possibility of its involvement in mediating tolerance in our carboplatin-resistant cell line.

### Pharmacological activation of Wnt signaling in wild-type cells disrupts carboplatin-response and enhances stemness and pluripotency marker expression

We hypothesized that modulation of Wnt signaling in the parental 468’P cell line could recapitulate the carboplatin-tolerant phenotype and increase the expression level of pluripotency and stem cell-related genes. To test this hypothesis, we treated 468’P with a small molecule inhibitor of GSK3β, CHIR99021 (CHIR), thereby preventing β-catenin degradation and consequently activating the Wnt/β-catenin pathway. To confirm Wnt signaling activation, we used a lentiviral fluorescent reporter of canonical Wnt transcriptional activity (TOPGFP) (48). 468’P displayed low basal levels of Wnt-reporter activity but promptly induced the reporter upon GSK3β inhibition, with almost 100% of cells becoming GFP positive within 12 hours of treatment (Fig. 2A).

**Figure 2.**
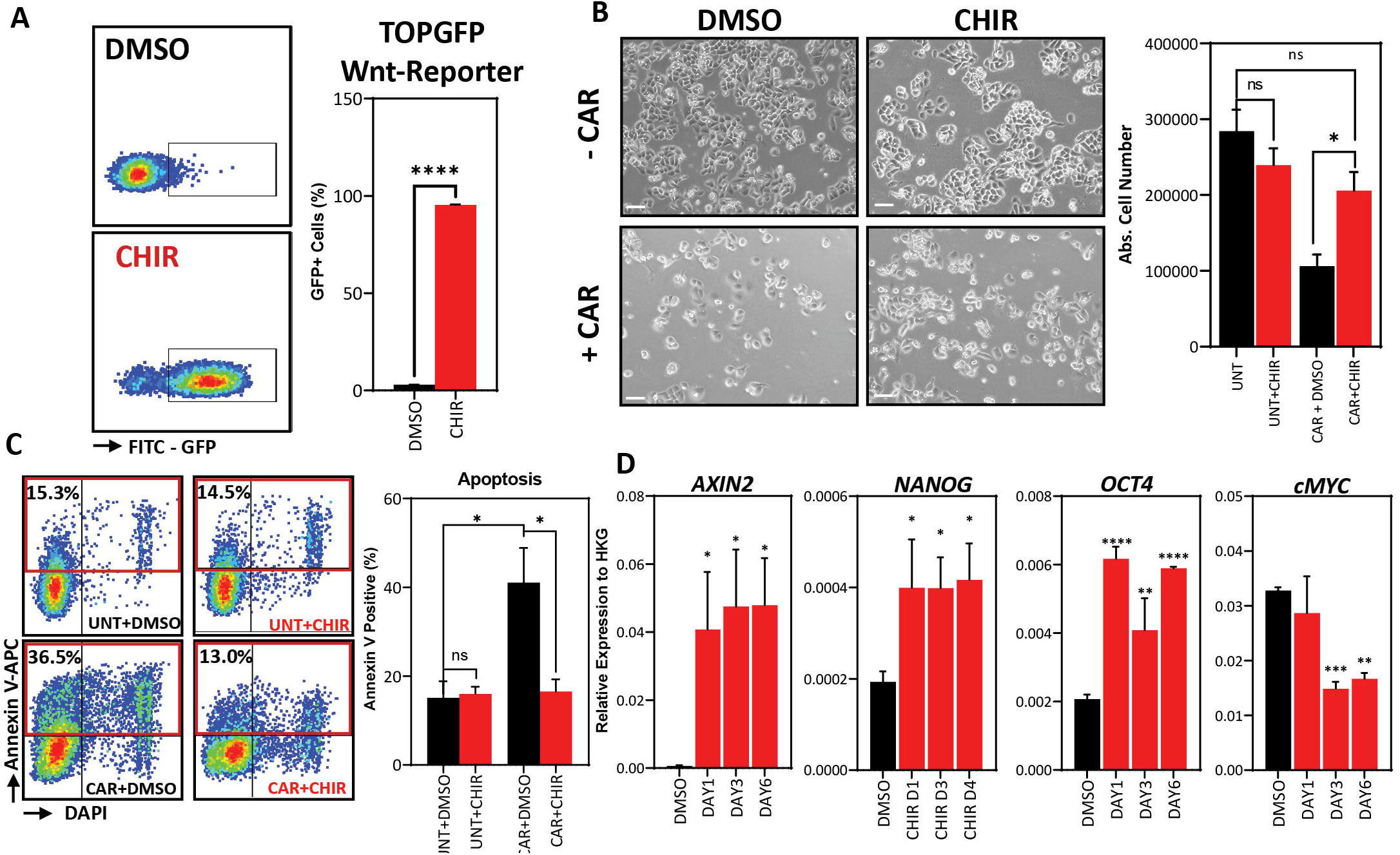
Pharmacological in vitro Wnt induction prevents carboplatin-induced apoptosis and upregulates stem cell marker expression. A) Representative flow cytometry scatterplots (left) of Wnt-reporter MDA-MB-468 TOPGFP cells treated with vehicle (DMSO) or GSK3β inhibitor (CHIR, 4 μM) for 12 hours and statistical analysis of the mean frequency of GFP+ cells using Welch’s t-test (right, n=3). B) Phase-contrast microscopy images of 468’P cells treated with or without carboplatin in the presence of CHIR or DMSO (left, scale bar: 100 μm) and statistical analysis of absolute cell numbers after 72 hours of each treatment using One-way ANOVA corrected for multiple comparisons using the Holm-Sidak method (n=4). C) Representative flow cytometry scatterplots of annexin V staining of 468’P cells (left) treated with or without carboplatin in the presence of DMSO or CHIR (4 μM) for 72 hours and statistical analysis of the mean frequency of annexin V positive cells (right) using one-way ANOVA corrected for multiple comparisons using the Holm-Sidak method (n=3). D) Relative mRNA expression of Wnt target and stem cell markers upon 72-hour treatment with DMSO or CHIR (4 μM) in 468’P cells (n=3). Multiple t-tests with Holm-Sidak correction for multiple comparisons. (Barplots represent mean + SEM. * *p <0.05, ** p<0.01, *** p<0.001, **** p<0,0001*, ns= non significant)

We observed a significant rescue of survival when treating 468’P with CHIR combined with carboplatin (Fig. 2B). Carboplatin-induced apoptosis was also significantly reduced when cells were co-treated with CHIR and CAR, as determined by flow cytometric analysis of Annexin V positivity (Fig. 2C). Also, GSK3β inhibition led to the upregulation of *OCT4* and *NANOG* pluripotency markers (Fig. 2D).

Similar results were observed by inhibiting GSK3β on a second TNBC cell line (MDA-MB-231) (Supplementary Fig.2 A-B), suggesting this effect is not cell line-specific.

### β-catenin overexpression induces carboplatin tolerance in 468’P and enhances stem cell features

GSK3β is a multi-substrate serine-threonine kinase that regulates a multitude of signaling pathways (49). As such, we proceeded to investigate whether the effects of pharmacological activation of Wnt signaling on stemness and carboplatin tolerance could also be induced by direct overexpression of β-catenin. To achieve this, we transduced 468’P cells with a lentiviral vector encoding a truncated, constitutively active mutant β-catenin (Δn90 β-catenin) (50), generating β-catenin overexpressing cells (468’OE) (Fig. 3A).

**Figure 3.**
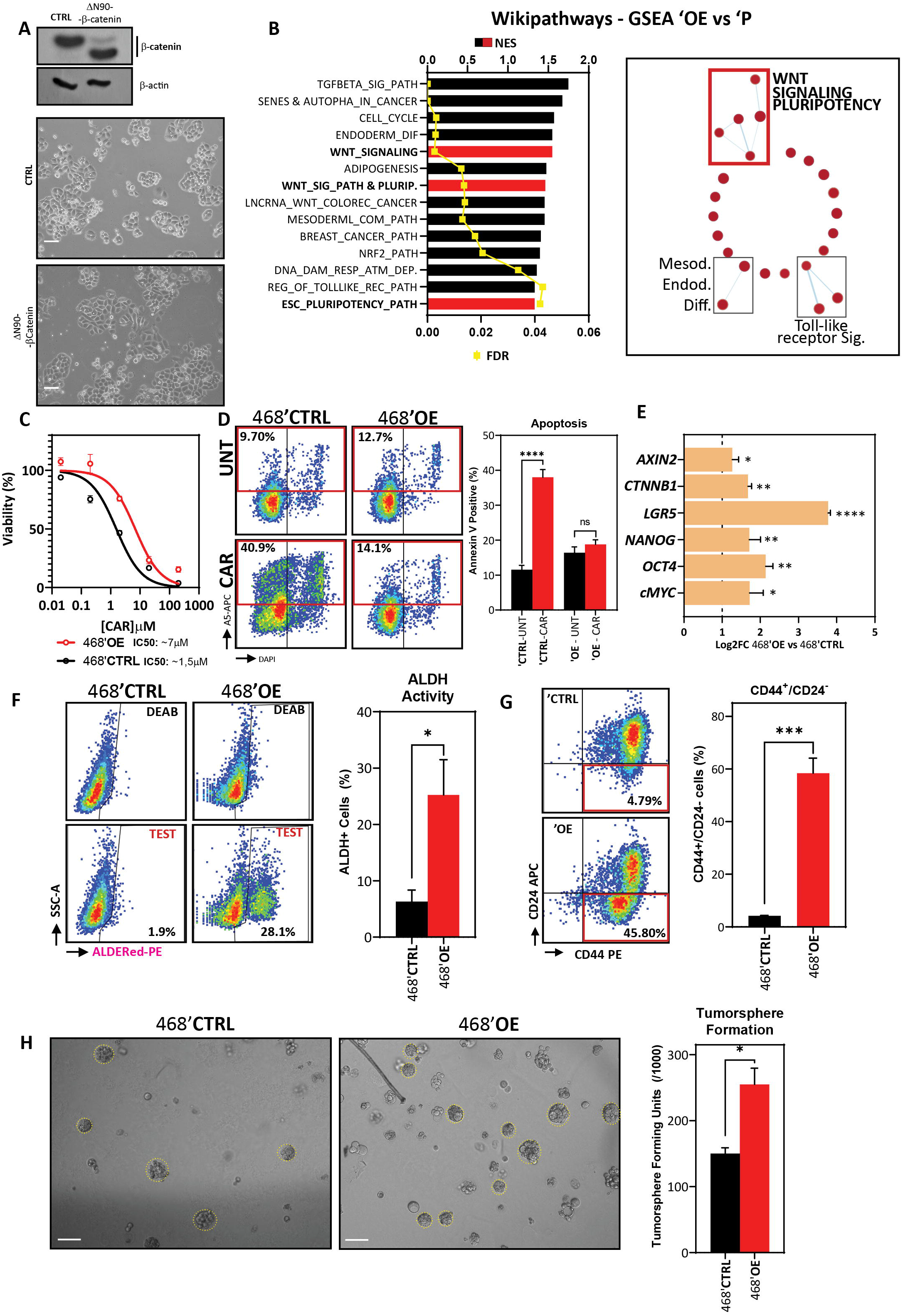
β-catenin overexpression in 468’P induces carboplatin-tolerance, pluripotency-related gene expression, and cancer stem cell features. A) Western blot (top) of total β-catenin in MDA-MB-468 cells transduced with an empty vector or truncated, constitutively active β-catenin isoform ΔN90 and phase-contrast microscopy (down). B) Enriched gene sets from Wikipathways database by one-tailed GSEA of ranked DEGs between 468’OE and 468’P sorted by normalized enrichment score (left) and enrichment map illustrating pathways most significantly different between 468’OE and 468’P (right). C) Non-linear fit model of [CAR] vs. normalized response for IC50 determination. (468’OE: n=6, R^2^=0.92; 468’CTRL: n=6, R^2^=0.95). D) Representative flow cytometry scatterplots of annexin V staining (left) of 468’CTRL and 468’OE cells treated with carboplatin 2 μM for 72h and statistical analysis of the mean frequency of annexin V positive cells using one-way ANOVA corrected for multiple comparisons using the Holm-Sidak method (right, n=3). E) mRNA level fold change (Log2) of *CTNNB1* (β-catenin), Wnt target *AXIN2*, and stem cell markers in 468’OE cells vs. 468’CTRL (n=4). Multiple t-tests with Holms-Sidak correction for multiple comparisons. F) Representative scatterplots of flow cytometric analysis of aldehyde dehydrogenase activity (left) and statistical analysis of the mean percentage of ALDH+ cells in 468’OE (n=5) and 468’CTRL (n=5) using Welch’s t-test (right). G) Representative scatterplots of flow cytometric analysis of CD44-PE and CD24-APC immunolabeling (left) and corresponding statistical analysis of the mean percentage of CD44^+^/CD24^−^ cells using Welch’s t-test (right; n=3). H) Representative brightfield images of tumorspheres generated from 468’CTRL and 468’OE cells (left, scale bar: 50 μm) and statistical analysis of mean tumorsphere forming units (number of spheres/number of seeded single cells) using Welch’s t-test (right; n=3). (Barplots represent mean + SEM. * *p <0.05, ** p<0.01, *** p<0.001, **** p<0,0001*, ns= non significant)

We then performed mRNA-sequencing to investigate biological processes altered by overexpression of β-catenin in TNBC cells. Differentially expressed genes (Supplementary Table 2) were ranked by fold-change and analyzed by GSEA using the Wikipathways database. We detected an interesting enrichment of gene sets related to both pluripotency and differentiation, namely Embryonic Stem Cell Pluripotency Pathways (M39530: WP_ESC_PLURIPOTENCY_PATHWAY) and Wnt signaling in pluripotency (M39387: WP_WNT_SIGNALING_PATHWAY_AND_PLURIPOTENCY) (Fig. 3B). In addition, GSEA revealed a significant correlation between the transcriptional profile of 468’OE cells and hESC and pluripotency transcription factor gene sets (Supplementary Fig. 3A).

468’OE displayed increased carboplatin tolerance in comparison with 468’CTRL cells (empty lentiviral vector) (IC50 468’OE: ~7μM vs. IC50 468’CTRL: ~1,5μM) (Fig. 3C). Accordingly, flow cytometric analysis revealed that upon overexpression of active β-catenin, 468’OE cells fail to induce apoptosis when challenged with carboplatin, displaying Annexin V positivity frequencies at the same level of untreated cells (Fig. 3D). These data confirm the direct involvement of β-catenin in the acquisition of *in vitro* carboplatin tolerance in TNBC.

Next, we probed gene expression of Wnt targets and pluripotency and cancer stem cell markers by RT-qPCR. As observed with pharmacological activation of Wnt, β-catenin overexpression also induced a significant upregulation of pluripotency markers *OCT4* and *NANOG* but also *cMYC* and *LGR5* (Fig. 3E).

Further, in accordance with the increased expression of pluripotency and stemness-related genes, flow cytometric analysis of ALDH activity (Fig. 3F) and CD44/CD24 expression (Fig. 3G) corroborated the presumptive induction of an enhanced cancer stem cell phenotype upon overexpression of β-catenin in TNBC cells.

Finally, we investigated whether overexpression of β-catenin functionally endows TNBC cells with enhanced tumorsphere formation capacity. Indeed, 468’OE cells displayed a significantly higher sphere-forming efficiency when grown in 3D suspension culture conditions, confirming that β-catenin overexpression functionally enhances *in vitro* stem cell properties in TNBC cell lines (Fig. 3H).

The overexpression of Δn90 β-catenin on MDA-MB-231 cells yielded the same effect with a roughly 10-fold increase in the IC50 of carboplatin, as well as increased ALDH activity and tumorsphere formation capacity (Supplementary Fig. 3 B-F).

### WNT inhibition disrupts carboplatin tolerance in 468’CT cells and downregulates cancer stem cell and pluripotency marker expression

Wnt signaling is deregulated in 468’CT cells, and β-catenin overexpression on parental cells confirmed its role in mediating the carboplatin-tolerant stem-like phenotype. Given these observations, we hypothesized that inhibition of Wnt signaling could restore sensitivity in 468’CT cells. To that end, we used LGK974 (LGK), a small molecule inhibitor of the endoplasmic reticulum palmitoyltransferase porcupine (PORCN). This enzyme is responsible for processing Wnt ligands for secretion, therefore mediating a crucial step of Wnt-dependent signaling (51).

Combinatorial treatment of 468’CT cells with 2 μM of CAR and LGK increased carboplatin-sensitivity in a dose-dependent manner (Fig. 4A). Interestingly, LGK treatment alone (200nM) was insufficient to induce apoptosis in both 468’CT and 468’P. However, when added to carboplatin, LGK974 induced strong annexin V positivity in 468’CT cells, indicating a rescue of carboplatin sensitivity in the tolerant cell line (Fig. 4B).

**Figure 4.**
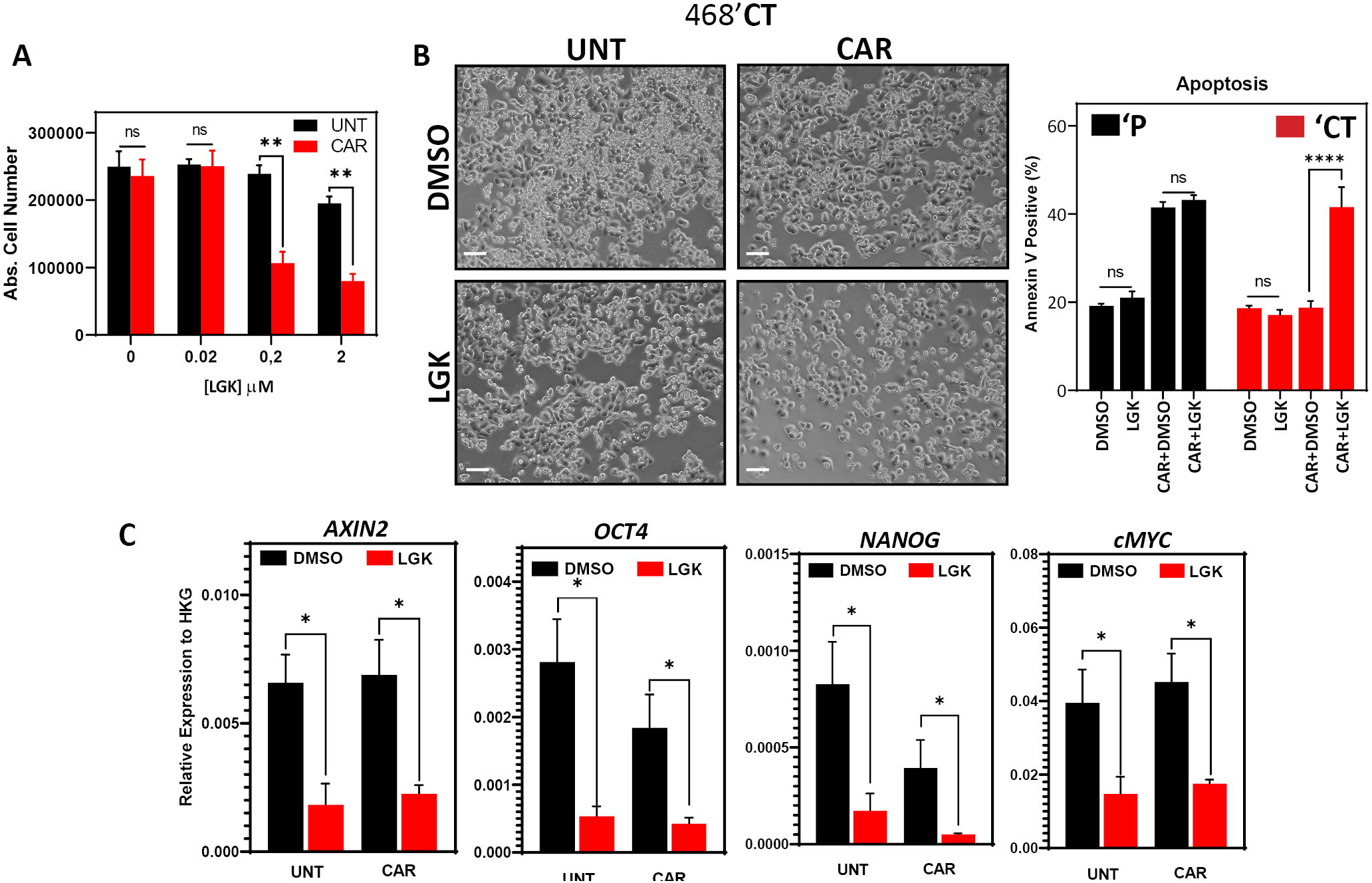
Wnt inhibitor LGK974 disrupts carboplatin resistance and pluripotency gene expression. A) Absolute cell number of 468’CT cells treated for 72 hours with or without LGK974 in the presence or absence of 2 μM carboplatin. Multiple t-tests (n=3). B) Phase-contrast microscopy of 468’CT cells treated for 72 hours with 200 nM LGK974 or DMSO in the presence or absence of 2 μM carboplatin (left). Mean frequency of annexin V positive cells in 468’P and 468’CT cells treated with or without 200 nM LGK974 in presence or absence of 2 μM carboplatin showing the resensitization of ’CT cells to carboplatin when co-treated with Wnt inhibitor (right, n=3). One-way Anova with correction for multiple comparisons using the Holm-Sidak method. C) Relative mRNA expression of Wnt target and stem cell markers upon 72-hour treatment with DMSO or 200 nM LGK974 with or without 2 μM carboplatin in 468’CT cells (n=3). Multiple t-tests: Unt vs. Unt+LGK & CAR vs. CAR+LGK. (Barplots represent mean + SEM. * *p <0.05, ** p<0.01, *** p<0.001, **** p<0,0001*, ns= non significant)

Gene expression analysis by qPCR of LGK974-treated 468’CT cells confirmed downregulation of Wnt signaling by reduced expression of WNT target *AXIN2*. More importantly, LGK974 severely reduced transcript levels of pluripotency markers *OCT4*, *NANOG*, and *cMYC* both in the presence or absence of carboplatin (Fig. 4C). These results corroborate the hypothesis that Wnt primes TNBC cells for carboplatin resistance by maintaining a stem-cell-like phenotype.

### Inducible β-catenin knockdown resensitizes 468’CT cells to carboplatin and disrupts expression of stem cell markers and tumorsphere formation capacity

Next, we investigated whether cancer stem cell markers and function could be manipulated by directly disrupting β-catenin making use of doxycycline (DOX) inducible short-hairpin RNA targeting *CTNNB1* (β-catenin) transcripts in 468’CT cells (iCTNNB1-KD) (52).

Gene expression analysis by RT-qPCR confirmed a reduction of roughly 90% in *CTNNB1* transcripts upon DOX treatment of iCTNNB1-KD cells whereas, as expected, cells expressing inducible scrambled shRNA (iSCRMBL) maintained basal β-catenin transcript levels (Fig. 5A). Notably, DOX-induced β-catenin knockdown led to a robust reduction of pluripotency and cancer stem cell markers, confirming the role of β-catenin in maintaining their high expression (Fig. 5A).

**Figure 5.**
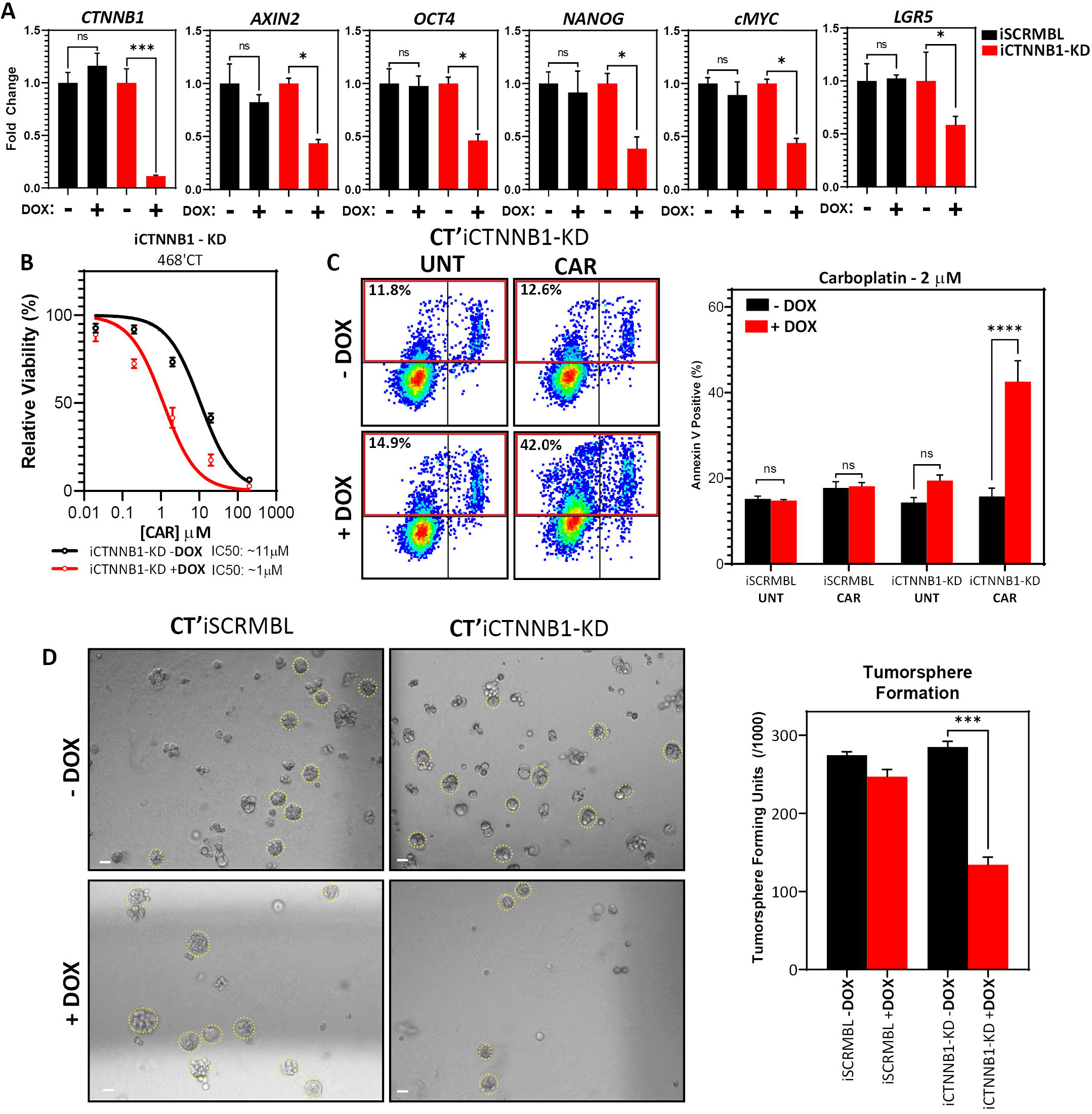
Inducible β-catenin shRNA disrupts carboplatin-resistance and stem cell function in 468’CT cells. A) Relative mRNA expression level of *CTNNB1* (β-catenin) and Wnt target and pluripotency markers in 468’CT cells transduced with inducible CTNNB1-targeting or SCRMBL shRNAs, in the presence or absence of doxycycline (n=3). Welch’s t-test (Dox vs. no Dox). B) Non-linear fit model of [CAR] vs normalized response for IC50 determination in iCTNNB1-KD cells in presence or absence of doxycyclin. (n=3, R^2^ + DOX: 0.93, R^2^ - DOX: 0.95). C) Representative scatterplots of flow cytometric analysis of apoptosis by annexin V staining of 468’CT iCTNNB1-KD and 468’CT iSCRBML cells treated with or without 2 μM carboplatin for 72h in presence or absence of doxycycline (left) and corresponding statistical analysis of the mean frequency of annexin V positive cells (right, n=6). One-way Anova with correction for multiple comparisons using the Holm-Sidak method. D) Representative brightfield images of tumorspheres generated from 468’CT iSCRMBL and 468’CT iCTNNB1-KD cells (left, scale bar: 50 μm) and statistical analysis of mean tumorsphere forming units (number of spheres/number of seeded single cells) (right; n=3). Welch’s t-test (Dox vs. no Dox). (Barplots represent mean + SEM. * *p <0.05, ** p<0.01, *** p<0.001, **** p<0,0001*, ns= non significant)

Next, we investigated the effect of β-catenin knockdown on *in vitro* carboplatin response by determining the IC50 for 468’CT iCTNNB1-KD cells in the presence or absence of DOX. Suppressing the expression of β-catenin had a remarkable effect on carboplatin tolerance. DOX-treated iCTNNB1-KD cells displayed a substantial reduction in measured IC50 compared to non-induced cells (IC50 - DOX: ~11 μM vs. IC50 +DOX: ~1 μM) (Fig. 5B). Importantly, 468’CT iSCRMBL cells displayed no significant changes in IC50 (-DOX: ~13 μM vs. +DOX: ~11 μM) (Supplementary Fig. 4). Additionally, when exposed to the IC50 of 468’P cells, 468’CT iCTNNB1-KD cells strongly induced apoptosis in the presence of DOX, while no difference was observed in iSCRMBL cells (Fig. 5C).

Finally, we assessed the effect of β-catenin suppression on tumorsphere formation as a functional readout for stem cell activity. In line with the downregulation of stem cell marker expression in the presence of DOX, iCTNNB1-KD cells displayed a significantly lower tumorsphere forming frequency when β-catenin shRNA was induced. On the other hand, differences upon the induction of SCRMBL shRNA were negligible (Fig. 5D).

### WNT inhibition disrupts *in vivo* carboplatin tolerance in a carboplatin-resistant TNBC Patient-Derived Xenograft

Patient-derived xenografts are essential *in vivo models* of human neoplasms. Moreover, PDX models retain with excellent fidelity histological and molecular features of originating tumors, therefore representing essential tools for assessing drug resistance and response (53–55).

To study whether, like in 468’CT cells, *in vivo* PORCN inhibition with LGK974 re-establishes carboplatin sensitivity, we used an isogenic carboplatin-resistant TNBC PDX (C4O) obtained from a previously chemotherapy-sensitive model (BRC016) (35). In brief, ten mice bearing BRC016 tumors were treated with 50 mg/kg of carboplatin once weekly for three weeks. Nine out of ten BRC016-bearing mice achieved a complete response to treatment with no tumors detectable. However, one tumor displayed a late response and, despite undergoing a substantial volume reduction, became tolerant to further treatment. The non-responder xenograft eventually regrew from the post-treatment residual tumor (Fig. 6A). Material from the regrown tumor was collected to establish an isogenic model of carboplatin-resistant TNBC (C4O) and gene expression analysis revealed changes in Wnt signaling target *AXIN2* and pluripotency and stem markers *NANOG*, *OCT4*, *SOX2*, and *LGR5* (Fig. 6B).

**Figure 6.**
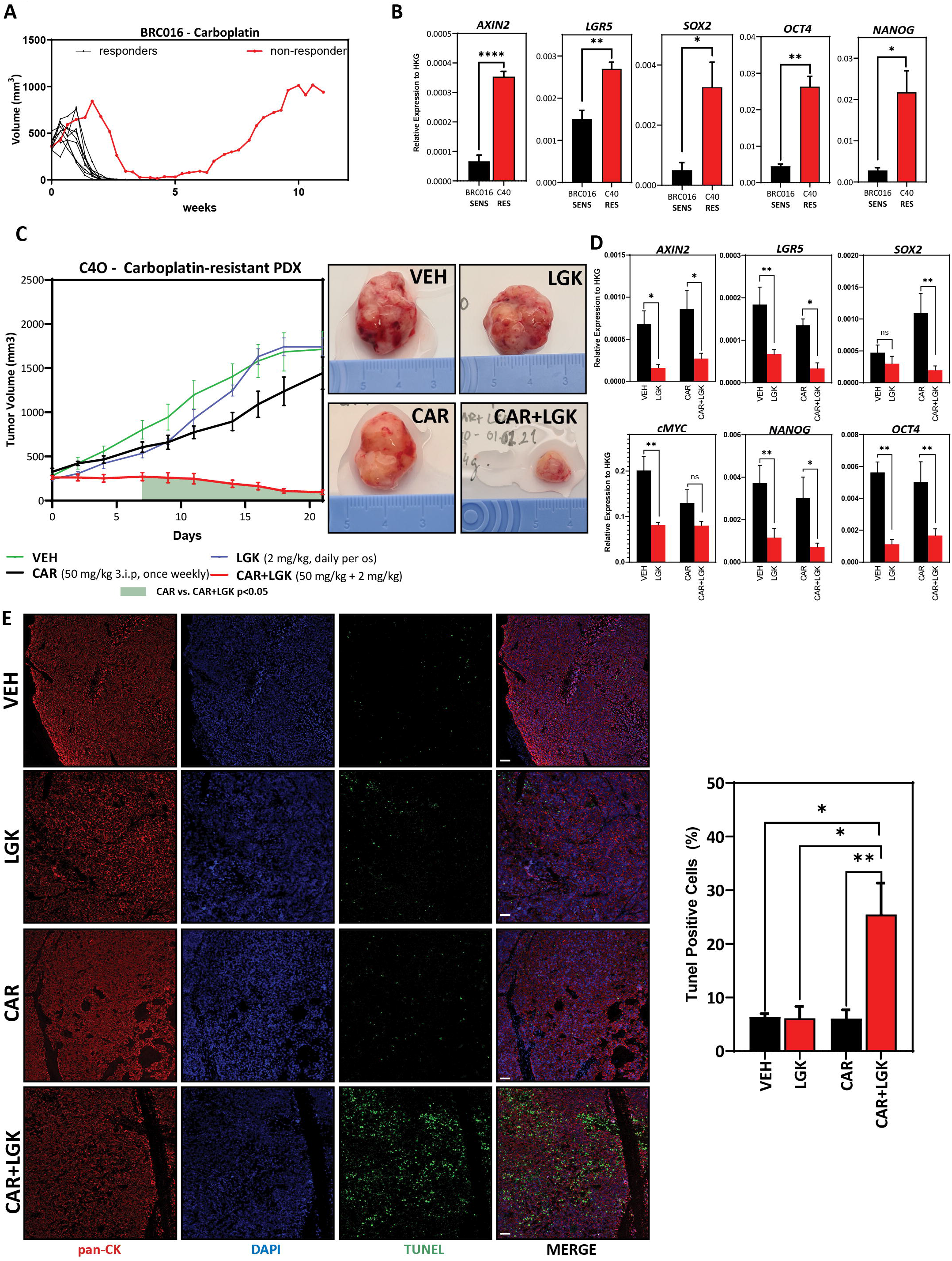
WNT inhibition disrupts in vivo carboplatin-tolerance in a carboplatin-resistant TNBC Patient-Derived Xenograft. A) Tumor growth curves of carboplatin-sensitive BRC016 TNBC PDX model. Nine out of 10 mice show complete response to treatment. One animal (red line) had a very delayed response and still had residual tumor mass after 3 weeks of treatment. The residual xenograft resumed growth after carboplatin-treatment was stopped. This tumor was collected to establish a carboplatin-resistant model (C4O). B) Comparative gene expression analysis by qRT-PCR of Wnt target AXIN2 and stem cell markers in BRC016 carboplatin-sensitive PDX and the C4O carboplatin-resistant isogenic PDX (n=4). Welch’s t-test. C) Tumor growth curves of C4O carboplatin-resistant PDX treated with VEH, LGK974, CAR, or CAR+LGK showing drastically reduced tumor growth in the combinatorial treatment arm (VEH, CAR, CAR+LGK n=6 and LGK n=5) (left). Two-way ANOVA with Tukey correction. The green-shadowed area under the curve represents highlights the time points in which the difference between CAR and CAR+LGK is statistically significant. Representative photographs of tumors in each treatment arm at day 21 of treatment (right). D) mRNA level fold change (Log2) vs. VEH treatment (no carboplatin and no Wnt inhibitor) in tumors dissected at treatment endpoint (21 days) (n=3). Multiple t-tests (VEH vs. LGK and CAR vs. CAR+LGK). E) Representative confocal microscopy images (left) of TUNEL staining in green and human pan-cytokeratin immunolabeling in red and respective quantification and statistical analysis of TUNEL positive cells (VEH n=4, LGK n=4, CAR & CAR+LGK n=6). One-way ANOVA with correction for multiple comparisons using the Holm-Sidak method. (Barplots represent mean + SEM. * *p <0.05, ** p<0.01, *** p<0.001, **** p<0,0001*), ns= non significant)

*In vivo* carboplatin tolerance was maintained in subsequent generations of transplanted C4O PDX models. We detected no significant differences in mean final tumor volumes in animals treated with vehicle or 50mg/kg carboplatin, administered once weekly intraperitoneally, for three weeks (Supplementary Fig. 5A). However, combinatorial treatment with daily dosing of PORCN inhibitor LGK974 drastically reduced C4O tumor growth (Fig. 6C and Supplementary Fig. 5A). Interestingly, LGK or CAR alone could not reduce C4O growth (Fig. 6C), and no significant differences in mean final tumor volumes between VEH, CAR, and LGK-treated animals were observed. Importantly, we did not find differences in Ki67 positivity between any treatment arms, indicating that reduced tumor growth in CAR+LGK treated animals was not due to differences in proliferation (Supplementary Fig. 5B).

Gene expression analysis by RT-PCR revealed that similar to what we observed upon treating 468’CT cells with LGK, inhibition of PORCN in C4O led to the depletion of pluripotency marker expression both in the presence or absence of carboplatin (Fig. 6D).

Finally, given the absence of alterations in the expression of Ki67, we sought out to understand whether the drastic reduction in tumor growth in animals treated with the combination of CAR and LGK could be due to increased apoptotic cell death. For this, we performed fluorescent terminal deoxynucleotidyl transferase dUTP nick end labeling (TUNEL) to detect DNA fragmentation as a readout of apoptosis. We combined TUNEL staining with human pan-cytokeratin immunolabeling to enable quantification of apoptotic signal specifically in cancer cells. No differences in TUNEL positivity were measured in VEH, LGK, and CAR treated animals. However, animals treated with the combination of CAR and LGK displayed a significant increase in apoptotic TUNEL signal (Fig. 6E)

### Wnt signaling is deregulated in patients with platinum-resistant TNBCs, high-grade serous ovarian cancer

To understand whether alterations in Wnt signaling are prevalent in platinum-resistant human TNBCs, we analyzed a public RNA microarray dataset (GSE103668) comprised of 21 pre-treatment samples from TNBC patients treated with cisplatin and bevacizumab (56). Due to the scarcity of additional platinum-treated TNBC transcriptomic data available in public repositories, we decided to include a supplementary dataset from high-grade serous ovarian cancer (HGSOC) (E-MTAB-7083) (Fig. 7A). The reason for this choice lies in the extensive use of platinum-based chemotherapy in this type of cancer and the striking overlap in clinical and molecular features between HGSOC and TNBC (57).

**Figure 7.**
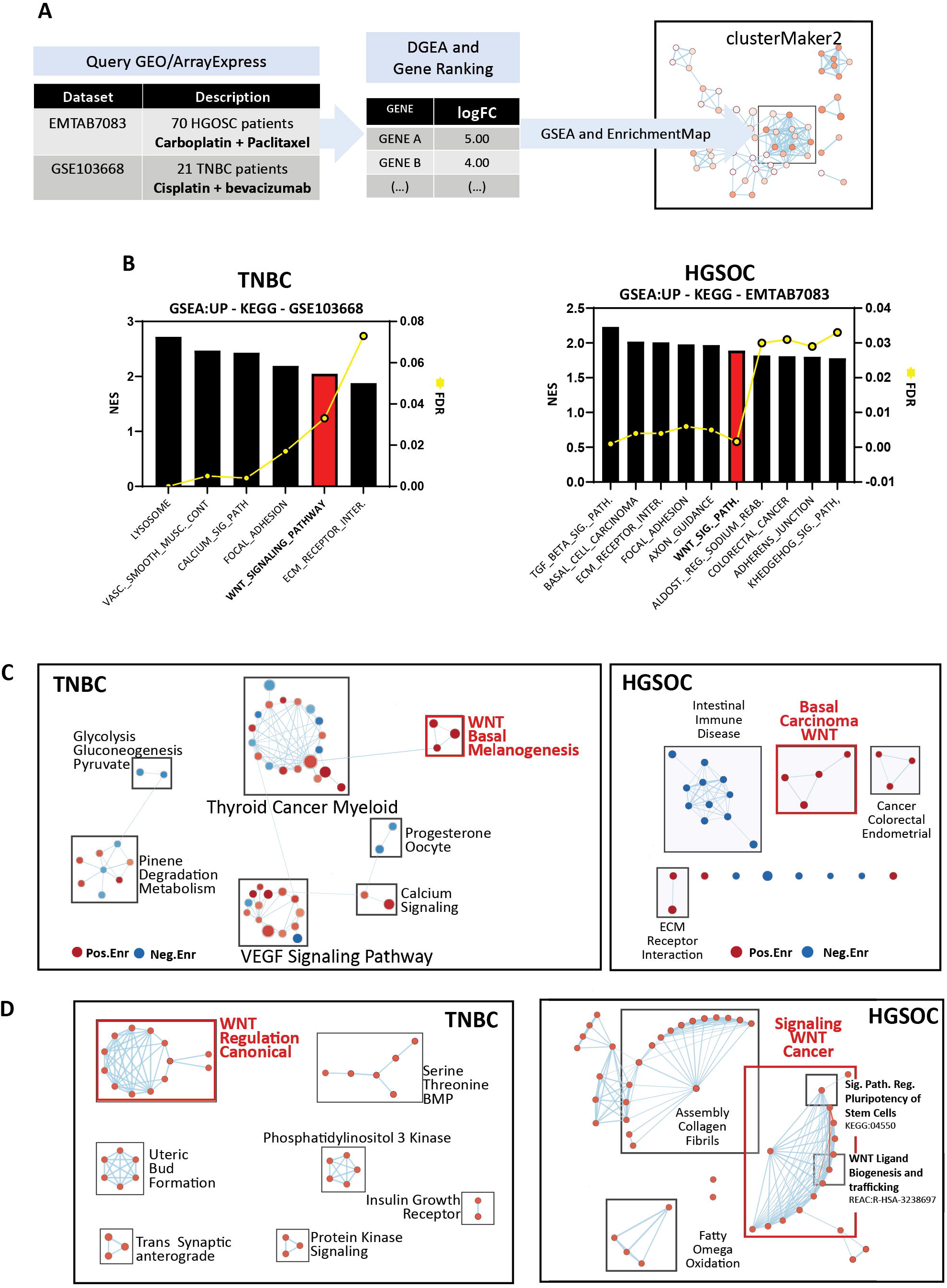
Wnt signaling is deregulated in patients with platinum-resistant TNBCs, high-grade serous ovarian cancer, and isogenic cisplatin-resistant ovarian cancer cell lines. A) Summary of datasets and analysis methodology used. Functional enrichment and mapping as previously reported (40,41,78). B) Enriched KEGG gene sets in patients with platinum-resistant TNBC (left) and HGSOC (right). C) Enrichment maps for visualization of enriched KEGG gene sets in patients with platinum-resistant TNBC (left) and HGSOC (right). D) Enrichment maps for visualization of gProfiler functional enrichment analysis of ranked, upregulated DEGs in patients with platinum-resistant TNBC (left) and HGSOC (right).

In both human datasets, the available clinical metadata was analyzed to classify patients based on response to treatment, and differential gene expression was calculated between responders and non-responders. Interestingly, GSEA analysis on ranked differentially expressed genes using the KEGG database retrieved Wnt Signaling as one of the top enriched terms with False Discovery Rate (FDR) <10% in TNBC and HGSOC patients with no response to platinum therapy (Fig. 7B). Other enriched KEGG terms comprised biological processes such as focal adhesion, extracellular matrix interaction, and other signaling pathways such as TGFβ and Hedgehog. Enrichment maps of GSEA hits for both datasets contained distinctive Wnt-related clusters involving gene sets with overlapping enriched genes such as “Melanogenesis” and “Basal Cell Carcinoma” for TNBC patients and HGSOC, with the addition of “Hedgehog Signaling Pathway” in the latter (Fig. 7C). To obtain a broader perspective of the function of differentially expressed genes, we performed functional enrichment analysis for both datasets using gProfiler to retrieve enriched gene ontology, Reactome, and Wikipathway gene sets to build enrichment maps. In both datasets, we obtained a distinctive cluster of Wnt-related terms, including regulating both canonical and non-canonical Wnt signaling in TNBC and regulating pluripotency and Wnt ligand biogenesis and secretion in HGSOC (Fig. 7D).

Altogether, these results highlight Wnt signaling’s importance in mediating platinum resistance in human TNBC and suggest transversal resistance mechanisms across TNBC and HGSOC.

## 4 Discussion

Primary and acquired resistance to chemotherapy poses a critical hurdle in the treatment of cancer. This is particularly important in TNBC due to the relatively limited therapeutic toolbox available and the daunting clinical characteristics of this disease. In the continued absence of targeted molecular therapies, we must strive to improve response to current therapeutic options given the high probability of shorter survival when pCR is not achieved. The use of platinum compounds in combination with other agents or as a standalone treatment in TNBC is still under intense investigation but already shows the potential to improve pCR rates in this breast cancer subtype. However, how TNBCs specifically develop resistance to platinum-based treatment is still a poorly understood process.

In this study, we used an isogenic carboplatin-resistant TNBC cell line and used next-generation mRNA sequencing to identify transcriptomic differences between sensitive and resistant cells. Functional enrichment analysis indicated, among others, the existence of profound differences in transcription of Wnt and pluripotency-related genes. We deemed this observation significant since Wnt signaling is intrinsically associated with tumorigenesis, and several studies have demonstrated the involvement of this pathway in mediating resistance to chemotherapy and radiation in different types of cancer, including breast (30,34,58–60). Wnt signaling is also known to specifically mediate platinum resistance in endometrial cancer (61), ovarian (62), and oral squamous cell carcinoma (63). Nonetheless, little is known regarding the role of Wnt signaling in mediating resistance to platinum in TNBC. Interestingly, Wnt is often deregulated in breast cancers, particularly TNBCs, despite the negligible frequency of mutations in Wnt pathway components. Significantly, Wnt activation is associated with poor clinical outcomes in TNBC (64).

Based on the transcriptomic data herein generated, we hypothesized that stem-cell gene expression and carboplatin resistance could be induced on parental MDA-MB-468 cells by manipulating Wnt signaling. For this, we first used CHIR, a small molecule inhibitor of GSK3β, thereby activating Wnt. Our results showed a significant increase in pluripotency marker expression and reduced apoptosis upon concomitant treatment with Wnt agonist and carboplatin. Past studies regarding the role of this multi-substrate kinase in treatment resistance are rather intriguing. While our results confirm other studies that report the enrichment of stem cell features upon GSK3β inhibition (65,66), they seem to contradict reports of GSK3 inhibition leading to reduction of tumor growth and apoptosis (67–69). Nonetheless, by overexpressing β-catenin in parental cells, we could replicate the phenotype of the isogenic carboplatin-resistant cells and GSK3β inhibition extensively. Namely, β-catenin overexpression induced a significant increase in expression of pluripotency markers and increased carboplatin tolerance, highlighting the role of Wnt pathway activation on drug resistance in TNBC.

Our results show an increase in the cancer stem cell population (CD44^+^/CD24^−^, ALDH^+^) concomitantly with higher expression of pluripotent markers such as *OCT4* and *NANOG* and *cMYC* and cancer stem marker *LGR5* in carboplatin tolerant model and Wnt active cells. LGR5 is known to maintain somatic and cancer stem cells in different tissues and cancers, including breast (46,70), to mediate cisplatin resistance in cervical cancer (71) and has been demonstrated to be a strong predictor of recurrence in estrogen receptor-negative breast cancer (72). Besides, NANOG expression predicts inadequate response to platinum in advanced non-small cell lung and oral squamous cell carcinomas (73,74).

Inhibition of Wnt ligand secretion using LGK974 in isogenic carboplatin-tolerant cells disrupted stem cell markers’ expression and reversed resistance. Interestingly, inhibition of Wnt ligand secretion alone was not enough to induce apoptosis in either carboplatin sensitive or resistant cells, despite severely downregulating stem cell markers’ expression. This indicates that Wnt secretory signals are not necessarily essential for the survival of either sensitive or resistant cells, but rather that these signals prime the latter for survival upon challenge by carboplatin.

LGK974 prevents the secretion of all Wnt ligands by inhibiting the palmitoyl acyltransferase PORCN. In vertebrates, the WNT family of lipid-modified secreted signaling proteins comprises 19 members, conferring a great deal of complexity to Wnt signaling. Wnt signaling includes a canonical or Wnt/β-catenin dependent pathway and the non-canonical or β-catenin-independent pathway (75). For that reason, it was essential to determine whether LGK974-induced carboplatin sensitivity was mediated directly by β-catenin. Silencing β-catenin downregulated stem cell markers’ expression and dramatically reversed tolerance to carboplatin, phenocopying LGK974 effects, and directly implicating canonical Wnt signaling.

Inhibition of Wnt ligand secretion, namely through the inhibition of PORCN, has been under scrutiny during the last decade as a potential therapeutic approach for different types of cancer. LGK974 specifically has shown excellent pre-clinical efficacy in Wnt-addicted models (76) and is under examination in phase I clinical trial for several Wnt-dependent solid malignancies, including TNBC (NCT01351103) (77). We evaluated whether this molecule could resensitize an *in vivo* model of carboplatin-resistant TNBC. Previous studies reported that LGK974 alone significantly reduces the growth of a murine mouse mammary tumor-Wnt3 model (76). To our surprise, daily dosing of LGK974 alone had no impact on tumor growth. This is analogous to what we observed when treating carboplatin-resistant cells with LGK974 alone. Importantly, we observed a significant reduction of tumor growth in animals treated with a combination of daily LGK974 and weekly carboplatin.

Finally, we were able to identify similarities in transcriptional profiles of patients with platinum-refractory ovarian and triple-negative breast neoplasms. Wnt signaling deregulation is a common denominator in both datasets we analyzed. Given the overlapping clinical and molecular features of both cancer types, it would be interesting to investigate whether transversal resistance mechanisms exist for other therapies.

Altogether, this study demonstrates that response to platinum can be improved and stable *in vitro*, and *in vivo* resistance can be reversed by a combinatorial approach to TNBC treatment which leverages the inhibition of Wnt-signaling to disrupt resistance-inducing cancer stem cell functions.

## Supporting information

Primer List

## 5 Data Availability Statement

The transcriptomic datasets generated in this study are available from ArrayExpress under the accession number E-MTAB-10337.

## 6 Ethics Statement

Tumor tissue for PDX implantation was acquired from a patient who provided written informed consent, and the procedure was approved by the Commission of Medical Ethics of the University Hospitals Leuven (approval numbers S54185 and ML8713). The PDX models herein used are now banked at the Trace Patient Derived Tumor Xenograft Platform of UZ Leuven/KU Leuven. All animal experiments were performed at the Trace PDX platform and were approved by the Ethics Committee Research UZ/KU Leuven (approval number P038/2015).

## 7 Author Contributions

WO and FL conceived and designed the study. SM generated *in-vitro* models of carboplatin resistance and the carboplatin-resistant C4O PDX under the supervision of DA and FA. YL generated the transgenic cell lines used in this study. WO and SM conducted all *in vivo* experimental work. EC conducted immunohistochemistry experiments. BV performed RNA-sequencing differential expression analysis, and WO performed downstream functional enrichment and visualizations. WO and partially YL carried out all flow-cytometry experiments. PA performed Western blots. WO carried out the remainder of the experimental work. Data analysis and figure preparation were performed by WO and reviewed by SM, DA, and FL. The manuscript was written by WO and reviewed, and approved by all authors. FL secured funding and supervised and guided experimental work and manuscript preparation. KK, JJV, MFB, and FA made comments on formatting, editing, and data analysis. All authors contributed to the article and approved the submitted version. WO and SM are co-first authors. DA and FLL are co-last authors.

## 10 Acknowledgments

We would like to give show their gratitude to Anchel de Jaime Soguero for his critical review of the manuscript. We also sincerely thank the Trace Leuven PDX platform for their help in setting up mouse experiments. Trace is part of the EurOPDX consortium.

## 11 Supplementary Material

Supplementary Table 1: Differentially expressed genes and GSEA of 468’CT vs 468’P

Supplementary Table 2: Differentially expressed genes and GSEA of 468’OE vs 468’P

Supplementary Table 3: Primer List

---

**Supplementary Figure 1.**
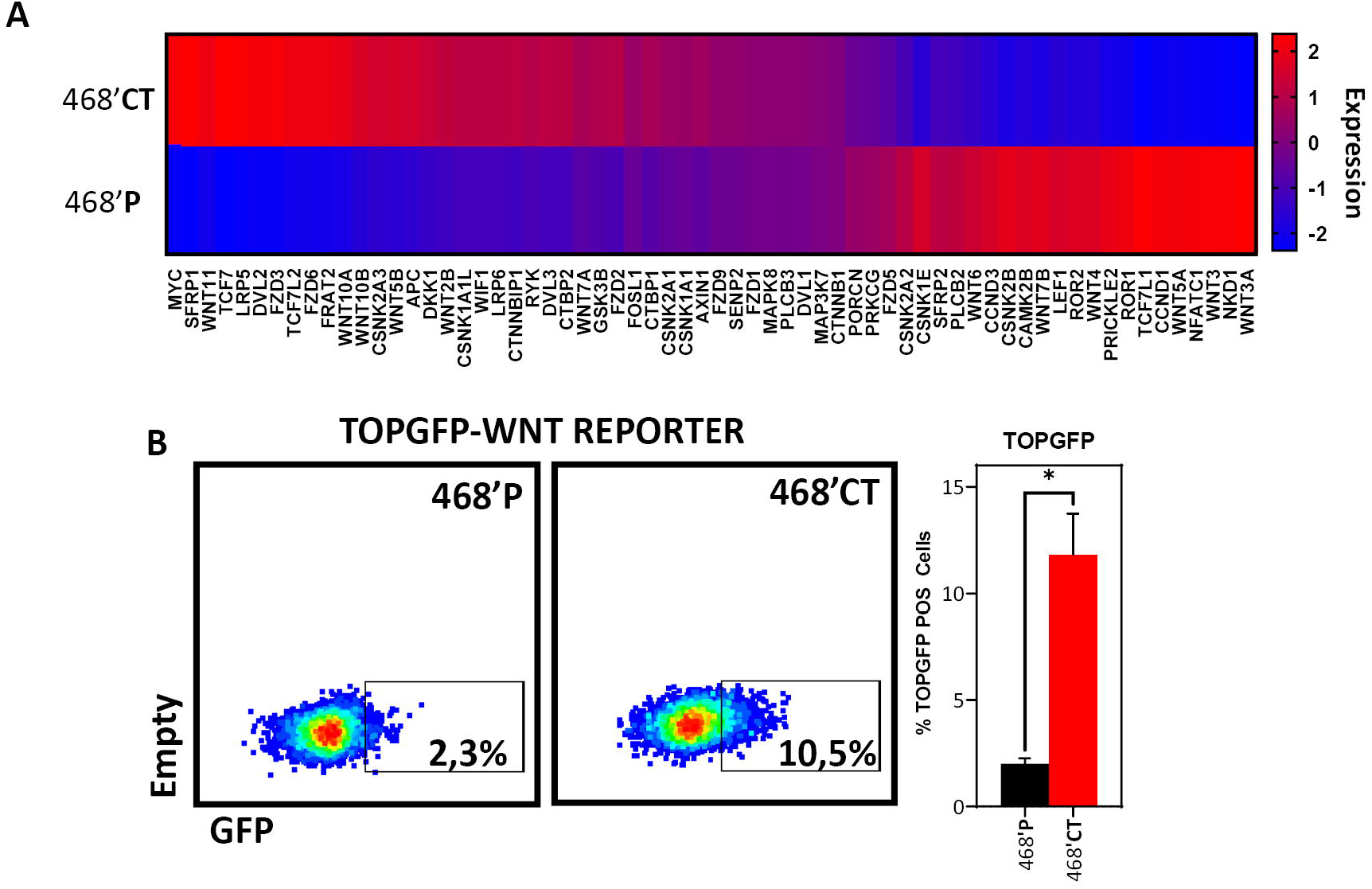
A) Expression heatmap of Wnt-related genes comprised in gene sets from the WNT Pathway Pluripotency cluster from Fig. 1E. B) Flow cytometry analysis of TOPGFP-Wnt reporter activity in 468’P and 468’CT cells (n=3). (Barplots represent mean + SEM. * *p <0.05, ** p<0.01, *** p<0.001, **** p<0,0001*, ns= non significant)

**Supplementary Figure 2.**
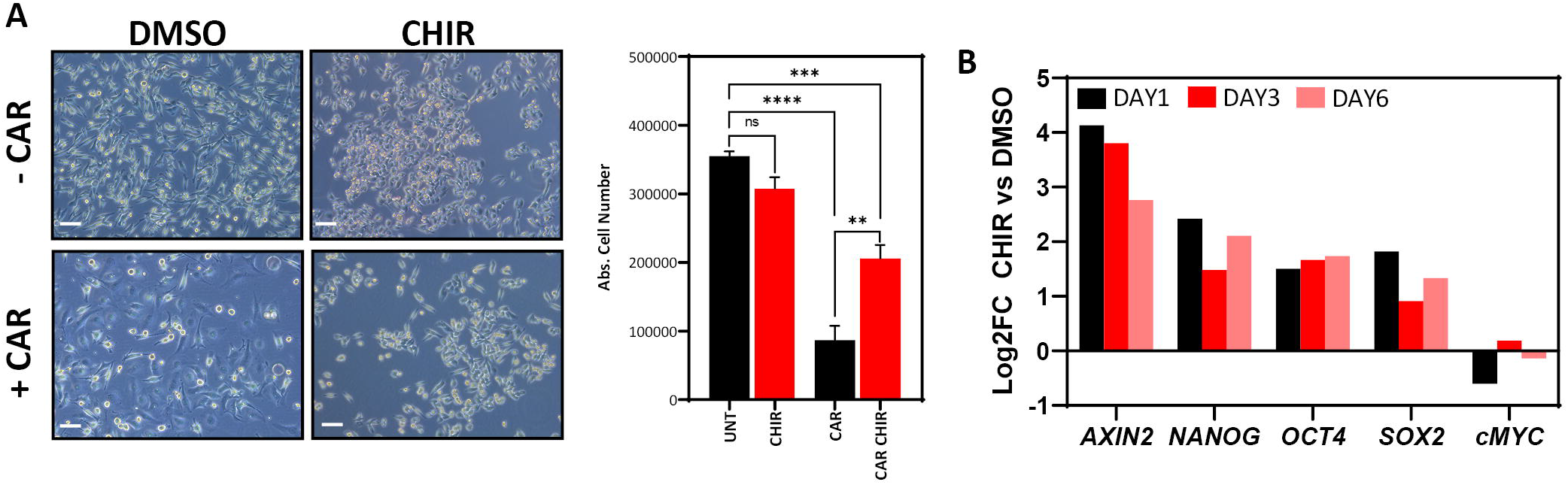
A) Phase-contrast microscopy (left, scale bar: 100 μm) of MDA-MB-231 cells treated for 72 hours with 35 μM Carboplatin in the presence or absence of 4 μM CHIR and statistical analysis of mean absolute cell numbers (n=4). One-way ANOVA with correction for multiple comparisons using the Holm-Sidak method. B) mRNA level fold change (Log2) Wnt target *AXIN2*, and stem cell markers in MDA-MB-231 cells treated with 4 μM CHIR vs. DMSO (n=1, average of technical replicates). (Barplots represent mean + SEM. * *p <0.05, ** p<0.01, *** p<0.001, **** p<0,0001*, ns= non significant)

**Supplementary Figure 3.**
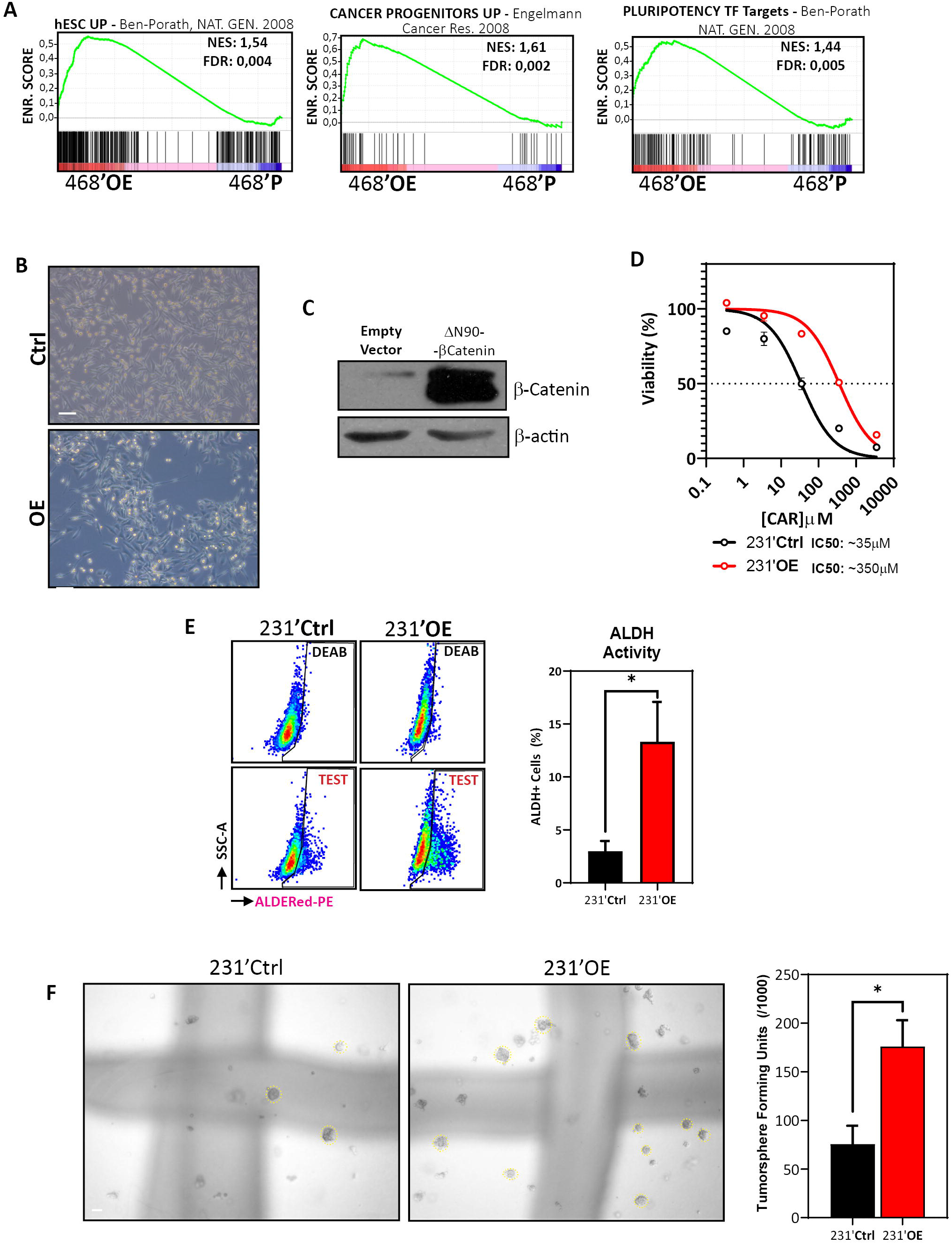
A) GSEA of hESCs (26) (left), Cancer Progenitor (44) (center), and pluripotency transcription target gene sets (26) (right) in 468’OE vs 468’P cells. B) Phase-contrast microscopy (scale bar: 100 μm) of control MDA-MB-23 and 231’OE (Δn90 β-catenin overexpression). C) Western blot of total β-catenin in MDA-MB-231 cells transduced with an empty vector or truncated, constitutively active β-catenin isoform ΔN90. β-actin was used as a loading control. D) Non-linear fit model of [CAR] vs. normalized response for IC50 determination (right). (n=2) E) Representative scatterplots of flow cytometric analysis of aldehyde dehydrogenase activity (left) and statistical analysis of the mean percentage of ALDH+ cells in 231’OE and 231’Ctrl cells using Welch’s t-test (n=5) (right). F) Representative brightfield images of tumorspheres generated from 231’Ctrl and 231’OE cells (left, scale bar: 100 μm) and statistical analysis of mean tumorsphere forming units (number of spheres/number of seeded single cells) (right; n=3). Welch’s t-test. (Barplots represent mean + SEM. * *p <0.05, ** p<0.01, *** p<0.001, **** p<0,0001*, ns= non significant)

**Supplementary Figure 4.**
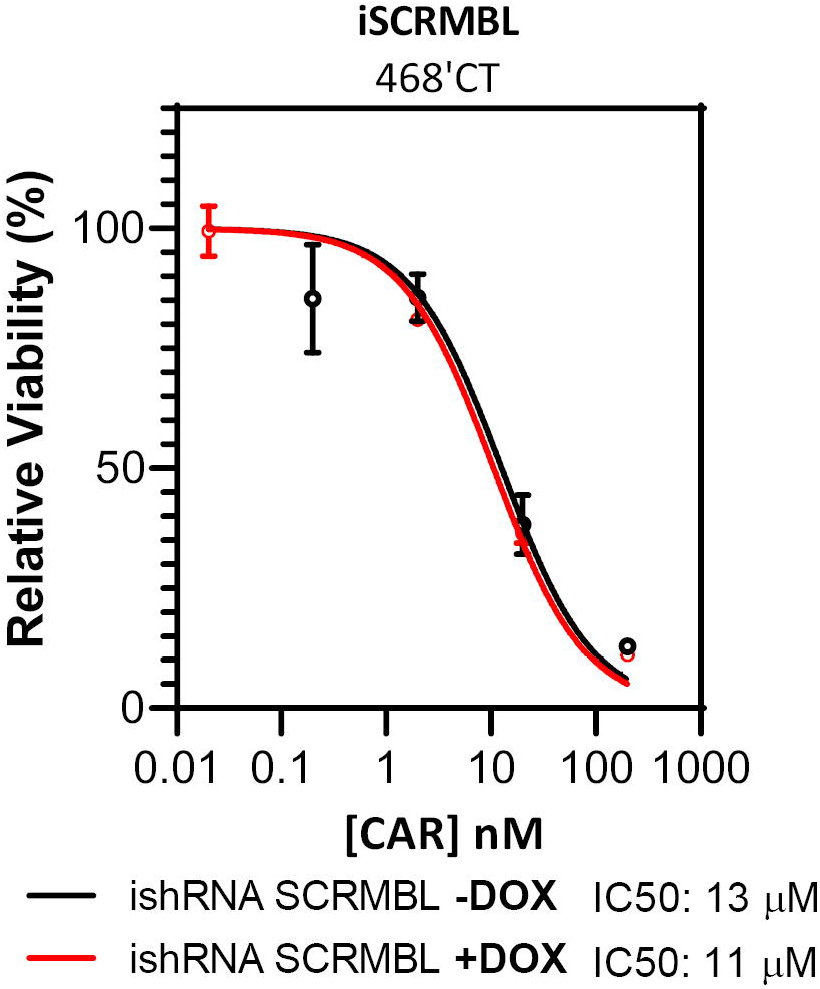
Non-linear fit model of [CAR] vs. normalized response for IC50 determination (right) in 468’CT cells transduced with inducible scrambled non-targeting shRNAs in the presence or absence of DOX. (n=2)

**Supplementary Figure 5.**
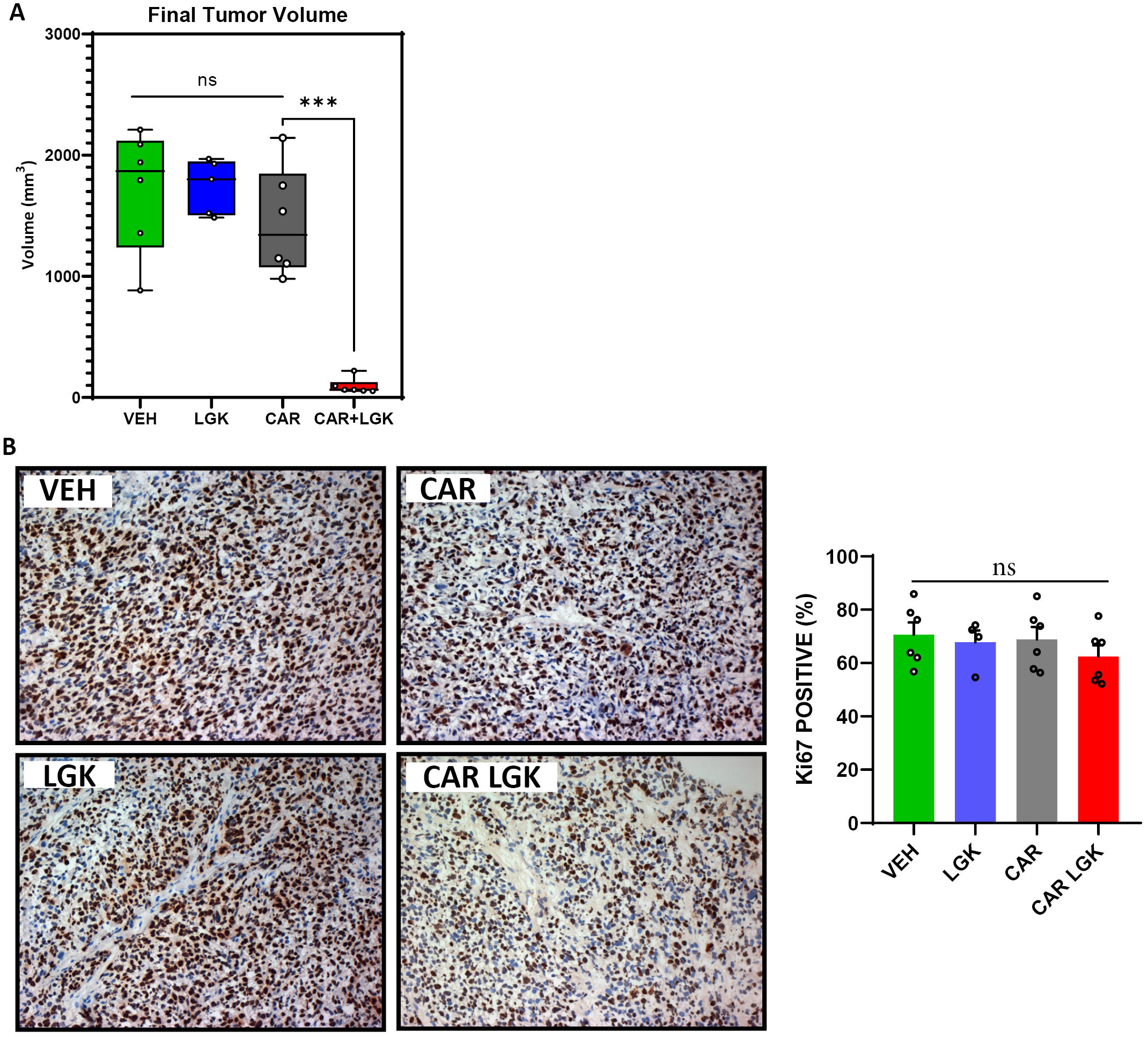
A) Final tumor volumes of VEH (n=6), LGK (n=5), CAR (n=6) and CAR+LGK (n=6) treated mice. One way-ANOVA with Holm-Sidak correction for multiple comparisons. B) Representative brightfield microscopy (20x magnification) of tumor sections labeled with anti-human Ki67 (left) and corresponding statistical analysis of the mean frequency of Ki67 positive cells. (VEH, CAR, CAR+LGK n=6 and LGK n=4). One-way ANOVA with Holm-Sidak correction for multiple comparisons. (Barplots represent mean + SEM. * *p <0.05, ** p<0.01, *** p<0.001, **** p<0,0001*, ns= non significant)

