## Supplementary material for "Wnt/β-catenin inhibition disrupts drug-tolerance in isogenic carboplatin-resistant models of Triple-Negative Breast Cancer": Primer List

| **Target** | **For/Rev** | **Sequence** |
| --- | --- | --- |
| AXIN 2 | FOR | AGTGCAAACTTTCGCCAACC |
|  | REV | TGAAGGACCTGTATCCACTGTC |
| cMYC | FOR | TGAGGAGACACCGCCCAC |
|  | REV | CAACATCGATTTCTTCCTCATCTTC |
| OCT4 | FOR | GTGGAGGAAGCTGACAACAA |
|  | REV | ATTCTCCAGGTTGCCTCTCA |
| NANOG | FOR | ACAACTGGCCGAAGAATAGCA |
|  | REV | GGTTCCCAGTCGGGTTCAC |
| SOX2 | FOR | TTGCTGCCTCTTTAAGACTAGGA |
|  | REV | CTGGGGCTCAAACTTCTCTC |
| RPL19 | FOR | AGTATGCTCAGGCTTCAGAAGA |
|  | REV | ATTGGTCTCATTGGGGTCTAAC |
| GAPDH | FOR | TCAAGAAGGTGGTGAAGCAGG |
|  | REV | ACCAGGAAATGAGCTTGACAAA |
| LGR5 | FOR | CTTACGTCACTGATGGTGCTTGC |
|  | REV | CTTGGAGAAAGAGATTTAGCCAGG |
| CTNNB1 | FOR | CCCACTAATGTCCAGCGTTT |
|  | REV | AACGCATGATAGCGTGTCTG |
